# The effects of predator recovery and climate change on the long-term demography of a flagship herbivore

**DOI:** 10.1101/2025.05.16.654556

**Authors:** Kenneth Loonam, Casey Brown, Darren Clark, Mary M. Rowland, Michael Wisdom, Taal Levi

## Abstract

In the western United States, ecosystems are being reshaped from both bottom-up and top-down processes. Widespread carnivore recolonizations after 20^th^ century extirpations have returned top-down pressures to ecosystems, and climate change is reshaping bottom-up forces through shifts in the timing and length of growing seasons, resulting in reduced forage quality and quantity. Large herbivores, such as elk (*Cervus canadensis*), function as flagship species for conservation and have potential for both top-down and bottom-up forces to shape demography as ecosystems change. To test the roles of top-down (puma density (*Puma concolor*)) and bottom-up (drought severity and elk density) processes in shaping large herbivore demography, and to tie those effects to population trajectories, we built an integrated population model (IPM) using 36 years of elk data spanning the recolonization of pumas and long-term climate change in eastern Oregon, USA. We tested the effects of top-down and bottom-up forces on elk recruitment, calf survival, and population growth. We also tested effects of bottom-up forces on elk pregnancy rates. Puma recolonization corresponded with declines in mean recruitment from 0.44 to 0.32 calves per female and in calf survival from 0.92 to 0.69, corresponding with a 4% drop in population growth. Drought severity increased over our study period, explaining a decline in mean recruitment from 0.39 to 0.32 calves per female, which corresponds to a 3% reduction in population growth. Drought severity likely acted on recruitment through reduced pregnancy rates. Lactating females showed lower pregnancy rates in drought conditions, while drought from fall breeding periods to the following fall had no effect on recruitment. While both top-down and bottom-up forces explained variation in elk demography over the course of our study, bottom-up forces are likely to be more influential on large herbivores moving forward. Predator populations stabilized following recovery, limiting the top-down contribution to annual variation in vital rates. Climate change-induced drought patterns, however, are accelerating during summer and fall throughout large areas of the interior western U.S. Thus, our results indicate the potential for increasingly strong bottom-up negative effects on herbivores whose performance depends on a narrow period of summer nutrition.

**Open Research Statement:** Data are not yet provided. Many of the data from Starkey Experimental Forest and Range are already publicly available, and we are currently working to post data for this manuscript on the Forest Service Research Data Archive (FSRDA) linked to the rest of the data from Starkey. Upon acceptance, all data and the relevant code will be archived the FSRDA. Currently, the code for the paper is publicly available on Github at https://github.com/keloonam/starkey_ipm

## INTRODUCTION

Large mammals function as flagship species for conservation across the world due to their visibility, high space needs, and economic value (Danell et al., 2006; Gordon et al., 2004; Organ et al., 2012; Sutherland, 2008). For flagship species to be effective conservation tools, we must accurately assess the demographic health of their populations as ecosystems change due to anthropogenic influences. Across North America, populations of many species of large mammals were locally extirpated by the early 20^th^ century (Mahoney, 2009). In the mid-20^th^ century, reintroductions and harvest regulations allowed the recovery of some large herbivores, most notably deer (*Odocoileus spp.*) and elk (*Cervus canadensis*), and by the late 20^th^ century, policy changes permitted large-carnivore predators to begin recolonizing historical ranges (Anderson & Lindzey, 2009; Linnell et al., 2001; Miller et al., 2013; Organ et al., 2012).

Along with predator recovery, climate change in the western United States has accelerated to produce earlier, longer, and hotter drought during summer and fall seasons, with shorter, milder winters and decreased snowpack (Bradford et al., 2020; Hicke et al., 2022). Increased temperatures and decreased precipitation with climate change during summer and fall have created conditions that substantially reduce forage availability for large herbivores (Brown et al., 2022), potentially decreasing fecundity, recruitment, and survival through bottom-up forces (Aikens et al., 2020; Middleton et al., 2013; Vose et al., 2016; Wato et al., 2016). Meanwhile, robust carnivore populations and communities can decrease survival and recruitment of large herbivores through top-down forces (Griffin et al., 2011; Hebblewhite et al., 2002; Hervieux et al., 2014). Large herbivores are thus caught in the middle of shifts in bottom-up effects from climate change and top-down effects from carnivore recovery. The relative roles of climate change and predator community structure in herbivore population dynamics are challenging to disentangle, because long-term demographic datasets of large herbivores that span both predator-recovery and climate change timelines are rare (Gaillard et al., 1998). As a result, most inquiries into the roles of top-down and bottom-up effects on large herbivores use spatial separation as a substitute for time series data, use indirect measures of demographic rates (e.g., repeated counts), or have relatively short time series data that may not show longer-term trends (Brodie et al., 2013; Clutton-Brock & Sheldon, 2010; Gaillard et al., 2000; Johnson et al., 2019).

Elk are a flagship large herbivore for land conservation in North America with ongoing debate over the relative influences of top-down and bottom-up forces on their demographic rates (Creel et al., 2011; Eberhardt et al., 2007; Vucetich et al., 2005; White et al., 2011). In large herbivores like elk, changes in adult female survival have large effects on population growth rates, but adult female survival is generally high and stable across years (Brodie et al., 2013; Gaillard et al., 1998). Recruitment, which is more variable than adult survival (though less influential on growth rates for a given change in the respective probabilities), often explains more of the variation in elk population growth rate (Gaillard et al., 2000; Lukacs et al., 2018; Raithel et al., 2007). Pumas, wolves, and bears (*Ursus spp.*) can all reduce recruitment through predation on juvenile elk, and wolves can additionally reduce adult female survival, supporting the hypothesis of top-down limitation of elk populations (Brodie et al., 2013; Lukacs et al., 2018; Proffitt et al., 2020). However, the effects of predation on population growth rates are less clear, because predation may result in compensatory declines in mortality from other causes (Brodie et al., 2013; Griffin et al., 2011; Hebblewhite et al., 2002). Bottom-up limitation of elk populations is most likely to be influenced by annual and seasonal variation in forage quality and quantity, which, when poor, can reduce the nutritional condition of adult females entering the breeding season, potentially resulting in lower pregnancy rates and recruitment (Cook et al., 2004; Johnson et al., 2013; Lukacs et al., 2018; Watkins et al., 2023). Bottom-up limitation of elk populations is also possible through density dependence as both pregnancy rates and calf survival have been better explained by population size than by annual conditions for high density populations (Singer et al., 1997; Stewart et al., 2005).

Although both top-down and bottom-up effects can be important drivers of elk population dynamics, their relative magnitude is still poorly understood. Further, the importance of late 20^th^ century predator population recovery and climate change in driving contemporary changes in elk demography is not well understood. Making such inference requires data that span a range of predator densities, extend across years with variable climatic and vegetative conditions, and account for variation in population density. Integrating those data to assess relative effect sizes on elk demographic rates and whether changes in demographic rates lead to changes in population growth rates will build a more complete picture of elk population dynamics in changing ecosystems.

We used 36 years of data (1988 to 2023) from Starkey Experimental Forest and Range (Starkey, Fig. 1, (Rowland et al., 1997)) in northeastern Oregon to test the relative importance of top-down forces, density dependence, and density-independent, bottom-up forces on elk demographic rates including pregnancy probability, recruitment, and calf survival. We also assessed whether changes in those demographic rates resulted in changes to elk population growth. Under the top-down hypothesis, puma density, which increased during the study timeframe as puma populations recovered throughout Oregon following a ban on hound hunting in 1994 (Oregon Department of Fish and Game, 2017), would negatively correlate with elk recruitment, juvenile survival, and population growth. Under the density dependence hypothesis elk density was predicted to negatively correlate with recruitment, survival, pregnancy rates, and population growth. Under the density-independent, bottom-up hypothesis, drought severity was predicted to negatively correlate with pregnancy rates in the year of the drought. For recruitment, drought in the current year represents the conditions experienced during pregnancy and rearing, while drought in the prior year represents the conditions when cows become pregnant. For survival, drought in the current year represents surviving through drought, while conditions in the prior year represent surviving another year after having experienced drought.

**Figure 1:**
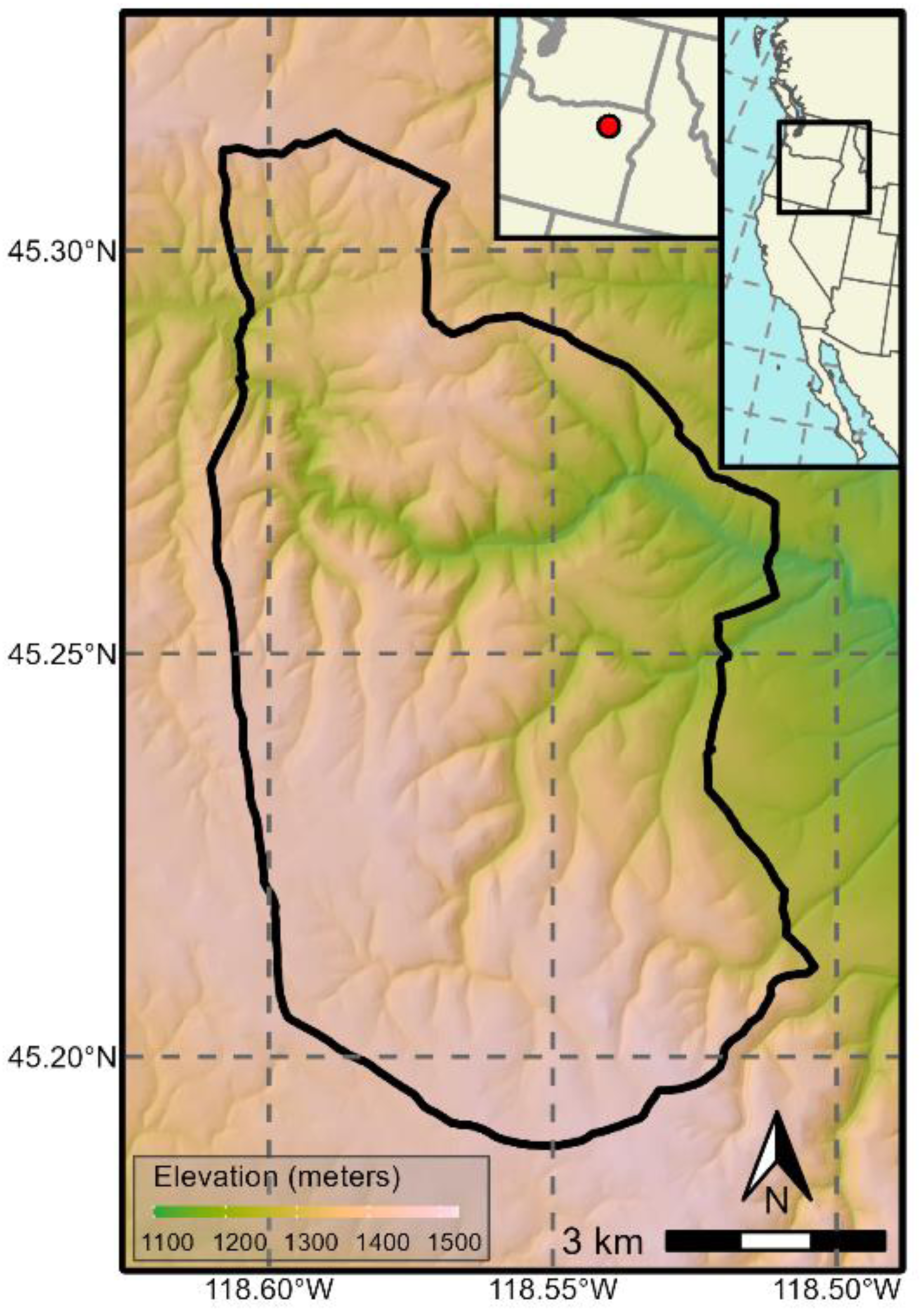
Primary study area within Starkey Experimental Forest and Range (black outline) with elevation represented by color. The inset maps show the location of Starkey in NE Oregon and the NW USA.

## METHODS

### Study Area

Starkey Experimental Forest and Range (Starkey) is in the Blue Mountains Ecoregion of NE Oregon and part of the Wallowa-Whitman National Forest (Fig 1). Starkey encompasses 100 km^2^ enclosed by a fence that is impermeable to elk but allows passage of smaller, non-ungulate species (Rowland et al., 1997) and carnivores (Ruprecht et al., 2021). Carnivores present include pumas, coyotes (*Canis latrans*), black bears, bobcats (*Lynx rufus*), and a relatively small, recent wolf presence, though none resided inside the fence on a regular basis during the study period (ODFW unpublished data, but see Ruprecht et al. 2021 for camera trapping methods used to collect the data). Elk are the most abundant wild ungulate. Mule deer (*Odocoileus hemionus*), white-tailed deer (*Odocoileus virginianus*), and cattle (*Bos taurus*) comprise the rest of large herbivore community, with cattle grazed seasonally from June through October under a standard federal grazing permit. The elevation in Starkey ranges from 1,122 to 1,500 meters, and the vegetative community is characterized by mixed coniferous forest interspersed with grass/shrubland.

### Elk Data Collection

The long-term elk monitoring in Starkey includes handling elk that voluntarily enter the winter feed grounds where they receive a maintenance diet (Starkey Experimental Forest and Range IACUC 92-F-0004; Wisdom et al. 1993). As part of handling, elk that arrive on the feed grounds are marked, building individual capture histories for each year (Wisdom et al., 1993). In addition to handling, age and sex classes are recorded including branch-antlered males, yearling males, antlerless adults, and calves. We combined yearling and branch-antlered males into a single, adult male, category resulting in three population counts: calves, adult males, and adult females. Adult females were further classified as young (< 3 years old), prime-aged (3-13), and old (>13) in the pregnancy analysis. Prior to being counted as calves in the fall, we refer to elk born each year as neonates. Elk harvested inside Starkey during controlled hunting seasons are reported and inspected by Starkey staff, as are non-harvest mortalities detected through monitoring collars, predation studies, or opportunistic observations. During inspections, individuals are aged and identified, which provides a terminal detection for previously captured individuals, novel IDs for previously unhandled individuals, and a census of harvested individuals broken down by sex and age class. Throughout Starkey’s history, a portion of the handled and harvested elk had lactation and pregnancy status recorded each year using blood samples to measure Pregnancy Specific Protein B (PSPB), or physical inspection of the uterus after harvest (Noyes et al., 1996, 2002). Females were recorded as lactating when milk could be extracted from the udder, which indicated that a female was still, or recently had been, nursing a calf. We used the pregnancy and lactation records of elk that had both statuses measured between November 1 and December 31 each year to ensure that lactation status accurately reflected whether the female nursed a neonate through the summer.

In addition to the harvest and feed ground monitoring efforts, Starkey researchers estimated elk abundance each year from 1989 to 2001 and in 2005 using aerial sightability and capture resight models (Unsworth, Kuck, and Garton 1990; Rowland et al. 1997; ODFW unpublished data). We incorporated the summary statistics from those models, including means and standard errors, as data in the integrated population model (IPM) (Eq. 11) (Schaub & Kéry, 2021).

### Bottom-up covariates

Starkey presents a unique opportunity to isolate the role of growing-season patterns as bottom-up influences on elk demography. Because most elk used Starkey’s winter area from December-March each year, a winter diet was provided during this period to minimize variation in animal condition associated with year-to-year changes in winter severity (Wisdom et al. 1993, Rowland et al. 1997). This approach allowed us to associate animal condition and population demography to the spring, summer, and fall periods (April through October), when variation in nutritional conditions and underlying weather patterns exert strong effects on animal condition in nearly all elk populations throughout the western U.S. (R. C. Cook et al., 2013). Additionally, though winter conditions have little effect on overwinter changes in elk condition in Oregon (Johnson et al., 2019), controlling for potential confounding effects through feeding allowed us to better isolate the effects of spring to fall conditions.

The extended time-frame of our study limits the available density-independent, bottom-up metrics. The gold standard, measuring the available digestible energy each year, was not possible. Instead, we tested multiple weather and remotely sensed variables, which are imperfect but effective proxies for forage conditions (Brown et al. 2022) that often correlate with elk demography (Brodie et al., 2013; Lukacs et al., 2018). Because we only had 35 observations of each demographic rate, we limited our tests to seven bottom-up covariates that represent the effects of variable growing seasons on elk demography (Table 3). We used the mean Palmer Drought Index (PDI) from April to September and the Standardized Precipitation and Evapotranspiration Index (SPEI) in September at three time-horizons (3, 6, and 12 months prior) as indices of precipitation and evapotranspiration during the growing season (Palmer, 1965; Vicente-Serrano et al., 2010), which both perform well at capturing local knowledge of drought conditions in the western U.S. (Heim et al., 2023). PDI is a cumulative measure of drought, e.g. the calculation for PDI includes the previous value of PDI and the temperature and precipitation that occurred between the two measurements. In contrast, SPEI measures the water balance using only the conditions during the specified time period. All three time-horizons for SPEI ended in September and represent the conditions elk experienced that year. The multiple time-horizons reflect hypotheses of the time scale on which water balance influences forage quality and elk nutrition in our system: the summer, full growing season, and full year for 3, 6, and 12 months, respectively (Table 2). In both PDI and SPEI, severe droughts (e.g. hot, dry conditions) are represented by negative numbers and above-average moisture conditions (e.g. cooler, wetter conditions) are represented by positive numbers, so the bottom-up hypothesis predicts a positive relationship between PDI/SPEI and calf survival, recruitment, pregnancy rates, and population growth. We also tested two simple weather variables, mean temperature and total precipitation from May through September (PRISM Climate Group, n.d.). Under the bottom-up hypothesis, calf survival, recruitment, pregnancy rates, and population growth will correlate with higher precipitation and lower temperatures. Finally, we tested the mean normalized difference vegetation index (NDVI) from May through September as a proxy for primary productivity, predicting it would positively correlate with elk demographic rates. We tested all of the density-independent, bottom-up covariates for correlations with recruitment and calf survival in the current year and at a one-year time-lag to disentangle the mechanisms through which they acted (Table 1). We only tested for current-year correlations with pregnancy rates, because we did not hypothesize any carryover effects in pregnancy probability (Table 1).

**Table 1:** Categories of covariates tested for correlations with recruitment, survival, and population growth in the IPM and pregnancy in a generalized linear model. Covariate category refers to the general mechanism being tested. For a list of the covariates tested in each category, see Table 3 and SI 1. The hypothesis/prediction column identifies the hypothesis each covariate tests and indicates the predicted direction of correlation under that hypothesis in parentheses.

| Covariate<br>Category | Hypothesis<br>(Prediction) | Mechanism |
| --- | --- | --- |
| Puma density | Top-down<br>(Negative) | *Recruitment*: predation on neonates in first 6 months post-birth (e.g. prior to counting) reduces calf/female ratios<br>*Survival*: predation on elk calves directly reduces survival |
| Elk density<sub>t-1</sub> | Density<br>dependence<br>(Negative) | *Recruitment*: competition for forage lowers body fat of adult females during fall breeding, lowering pregnancy rates and ability to invest in neonates if pregnant<br>*Survival*: competition for forage among female elk limits investment in nursing and limits calf growth going into winter |
| Drought<sub>t</sub> | Bottom-up –<br>density independent<br>(Positive) | *Recruitment*: cooler, wetter conditions improve forage, allowing a higher proportion of females to successfully nurse neonates over the summer.<br>*Survival*: cooler wetter conditions throughout the year improve post-weaning, calf forage, allowing higher survival |
| Drought<sub>t-1</sub> | Bottom-up –<br>density independent<br>(Positive) | *Recruitment*: cooler, wetter conditions improve forage and cow body fat, allowing for higher conception rates during fall breeding and greater investment in neonates.<br>*Survival*: cooler wetter conditions improve forage, allowing cows to invest more in neonates, which then enter the calf year with more body fat, increasing the odds of surviving the year |

**Table 2:** List of the time periods referenced in the text with the months that each includes. The IPM estimates abundance on November 1 of each year. Months subscript as *t* immediately precede November_t_.

| Season | Months | Notes |
| --- | --- | --- |
| Breeding | September<sub>$t$</sub> – October<sub>$t$</sub> | Estrus and breeding – when females become pregnant |
| Growing | April<sub>$t$</sub> – September<sub>$t$</sub> | Start of the spring green-up to the beginning of fall |
| 3-month SPEI | July<sub>$t$</sub> – September<sub>$t$</sub> | Peak to late summer – typically the hottest and driest months |
| 6-month SPEI | April<sub>$t$</sub> – September<sub>$t$</sub> | Aligns with growing season |
| 12-month SPEI | October<sub>$t-1$</sub> – September<sub>$t$</sub> | Full-year water balance leading up to mating |

**Table 3:**
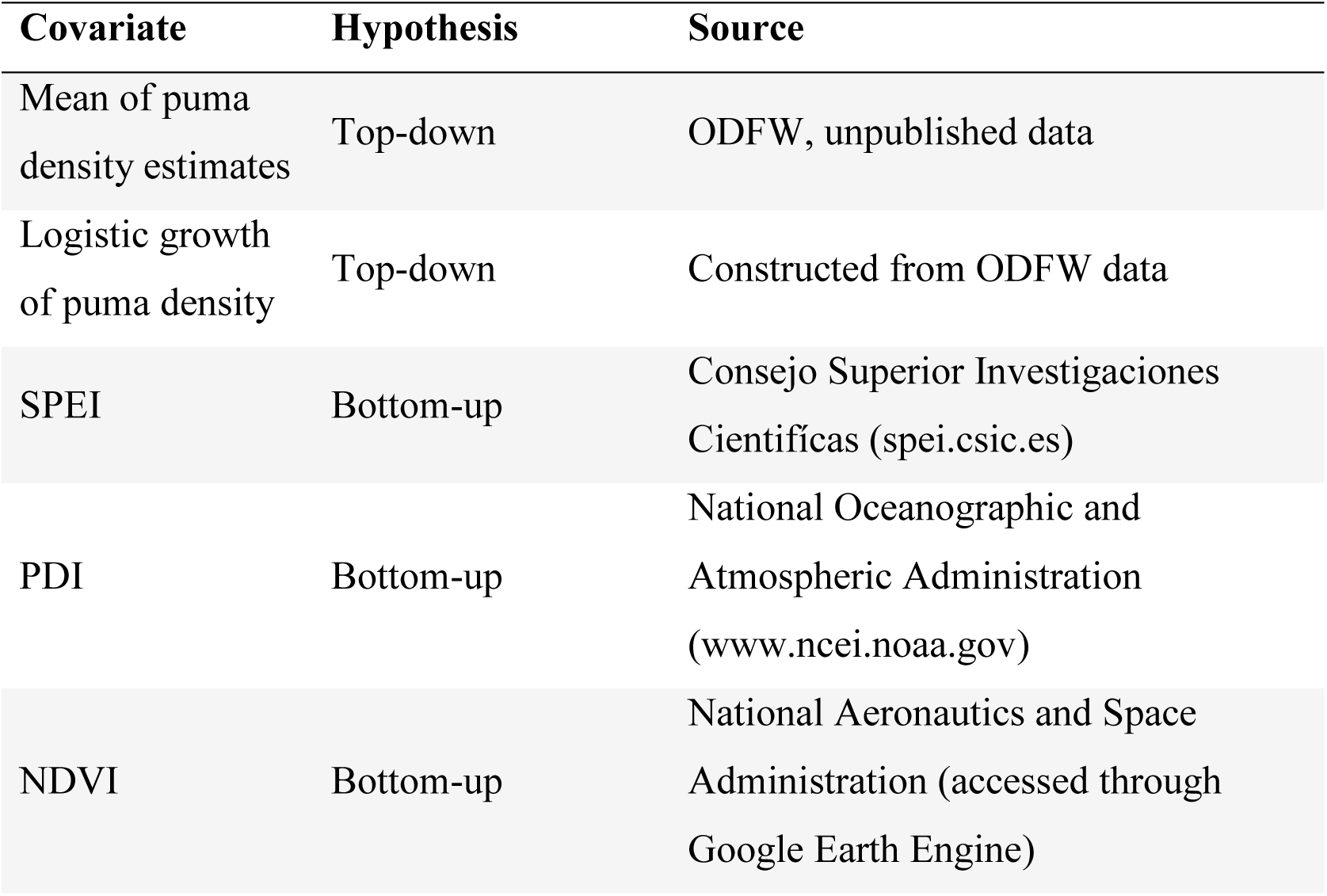
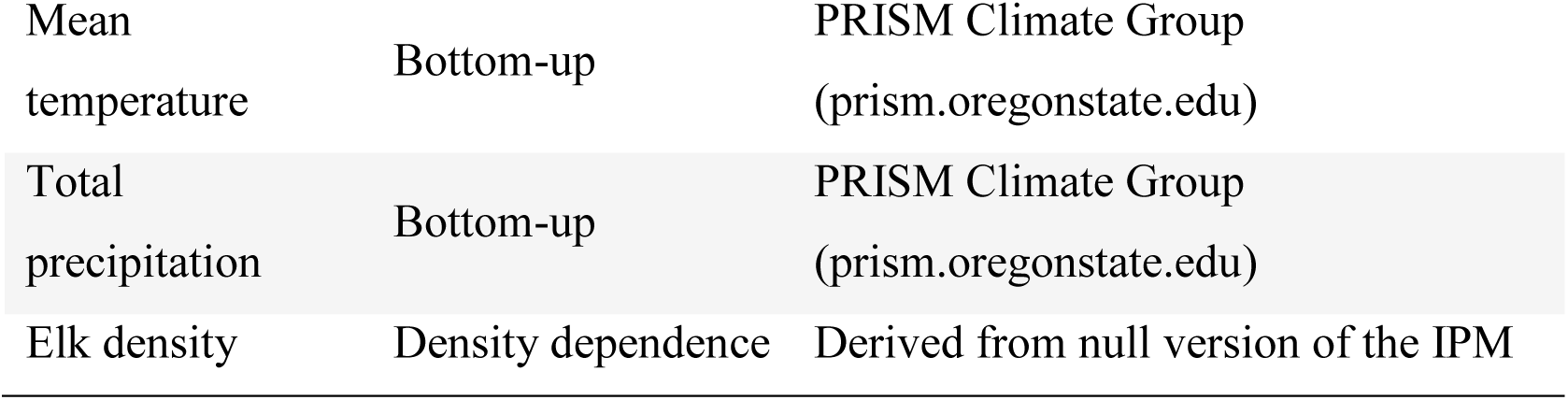
List of all covariates tested with descriptions and sources for each. We tested all of the drought metrics in models that included the covariate without a time-lag and with a one-year time-lag, e.g., a single model included two versions of the same covariate and each covariate was only tested in one model. For details on measurement timing relative to the process model, see Fig. 1 and Table 2. The mean of the puma density estimates is the arithmetic mean of two scaled estimates of puma density at varying spatial scales (SI 1). SPEI, the standardized precipitation evapotranspiration index, is a measure of the water balance in an area over a specified time-frame. We tested 3 time-frames, 3, 6, and 12 months, all ending in September (Table 2). PDI, the Palmer Drought Severity Index, is a cumulative measure of drought severity that incorporates the previous measure of PDI and the temperature and rainfall since the previous measure. For PDI, NDVI, and temperature, we used the mean value from April through September. The main text focuses on the mean estimate for puma density and on the 12-month SPEI for drought severity. For further discussion of models run, see SI 1.

| Covariate | Hypothesis | Source |
| --- | --- | --- |
| Mean of puma density estimates | Top-down | ODFW, unpublished data |
| Logistic growth of puma density | Top-down | Constructed from ODFW data |
| SPEI | Bottom-up | Consejo Superior Investigaciones Científicas (spei.csic.es) |
| PDI | Bottom-up | National Oceanographic and Atmospheric Administration (www.ncei.noaa.gov) |
| NDVI | Bottom-up | National Aeronautics and Space Administration (accessed through Google Earth Engine) |
| Mean<br>temperature | Bottom-up | PRISM Climate Group<br>(prism.oregonstate.edu) |
| Total<br>precipitation | Bottom-up | PRISM Climate Group<br>(prism.oregonstate.edu) |
| Elk density | Density dependence | Derived from null version of the IPM |

We also used female density from the prior year (i.e., the density females experienced leading up to and during the breeding season Fig 2.) as a covariate to test for bottom-up limitation through density dependence on recruitment, calf survival, population growth, and pregnancy. Including a direct relationship between recruitment and abundance in the model, however, creates a circular dependency. To avoid that, we ran a null version of the IPM and used the median posterior elk density from the previous fall as the elk abundance covariate. We compared the covariate estimate of density and the posterior from the full version of the IPM and found no difference (SI 1).

**Figure 2:**
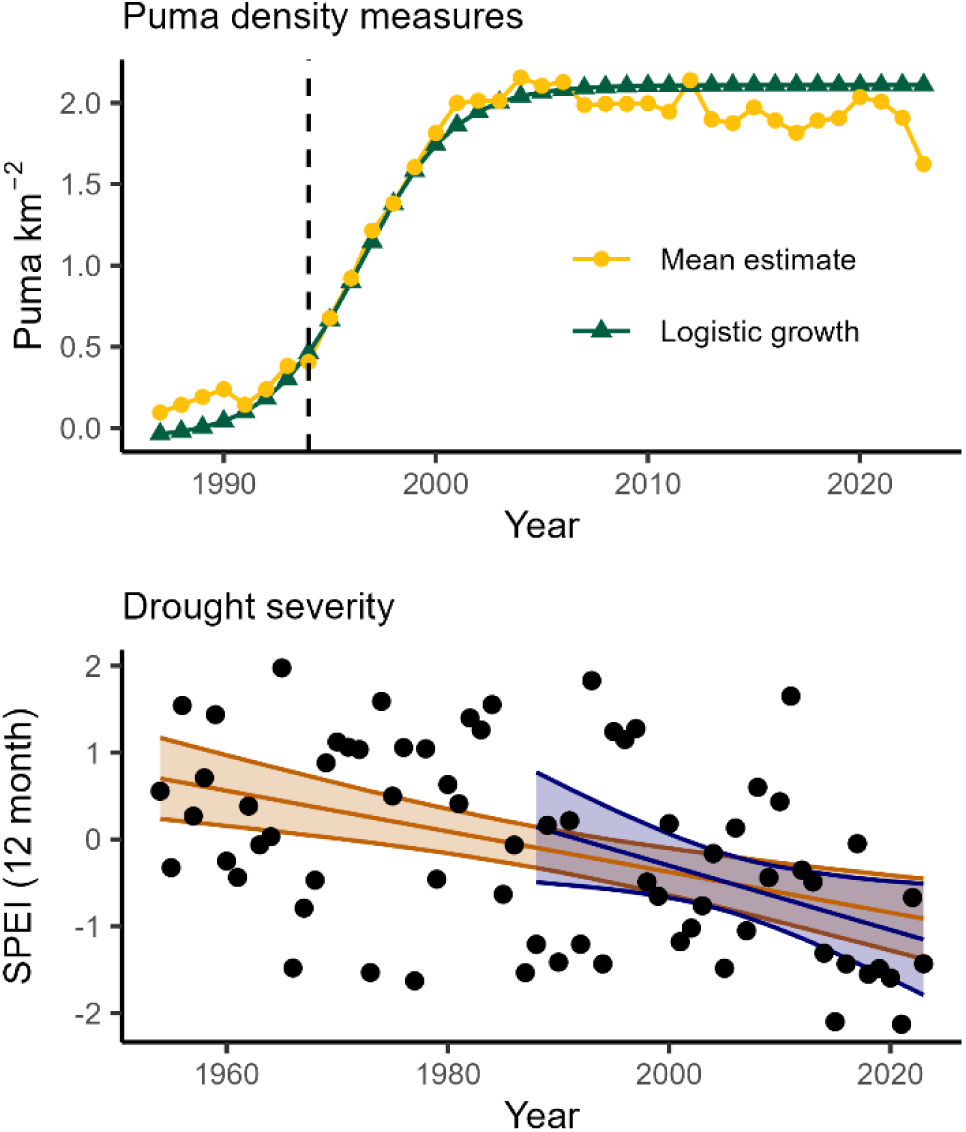
Annual values for the top-down covariates (top panel) and the most influential bottom-up covariate (bottom panel). In the puma density graph, the green line is a logistic growth curve fit to the Starkey WMU population reconstruction data (SI 1). The yellow line is the mean of the population reconstruction and an ODFW estimate of puma abundance for the entire Blue Mountains region. Prior to taking the mean, they were scaled using the years that they follow the same trend. In the drought severity graph, each point represents and observation of 12-month SPEI in September of a given year. The orange line is a linear best fit line over the entire time-span of the data (slope = −0.023; −0.035 – −0.012; P(d) > 0.99) and the orange shaded area represents the 95% credible intervals of the trend. The blue line and shaded area is the same regression but using only data from the time-frame of our study (1988-2023) (slope = −0.037; −0.068 – −0.005; P(d) > 0.99).

### Top-down covariates

To test the top-down hypothesis, we used a time series of puma density that spans the full period of the study (1988-2023). Under the top-down hypothesis, calf survival and recruitment will negatively correlate with puma density. Calf survival includes a year of predation risk (November to November), and recruitment includes 5 months of predation risk (neonates surviving from birth to November counts).

We constructed the time series of puma density from two sources (SI 1). First, we used a population reconstruction based on pumas harvested in the Starkey Wildlife Management Unit, which encompasses Starkey (Lancia et al., 2005; Stoner et al., 2006). The population reconstruction sums the number individuals in the population based on the years each individual was alive using age-at-harvest data. Because it relies on detections at death, recent years show an artificially declining population until new mortalities “fill in” the population in previous years. Because the population reconstruction is not reliable for recent years, we supplemented it with a puma population estimate of the Blue Mountains (which encompass Starkey WMU) from mortality and hunter success data in conjunction with a cougar population process model (Keister & Van Dyke, 2002) (SI 1). The Blue Mountain estimate and the Starkey WMU reconstruction show the same patterns until 2012: an initially small population that grew from 1993 to 2002 and was stable from 2003 to 2012 (D. Clark, ODFW, unpublished data). After 2012, the Blue Mountain estimate shows another growth period, and the Starkey WMU reconstruction shows a largely artifactual decline.

We took two approaches to combine the two sources of puma density into continuous indices of the puma population. First, we scaled each estimate using the mean and standard deviation of the years during which they showed the same patterns (1994-2012) and took the mean (SI 1), which we refer to as the mean of puma density estimates (Table 3). That puma density covariate shows logistic-shaped growth stabilizing in 2004, after which it fluctuates slightly below the peak (Fig. 2). Second, we used the population reconstruction from the Starkey WMU (the more fine-scale of the estimates), to fit a smoothed logistic growth curve (SI 1). To do that, we truncated the reconstruction at the year it begins declining (2013) to fit the curve, and projected it through 2023 using the estimated logistic parameters. The logistic growth version of puma density matches the mean of the estimates closely, but stabilizes at a slightly higher density and smooths over the inter-annual variation that is likely artifactual (Fig. 2). Both trends conform with the timing of puma recolonization, puma population growth following the ban on hound hunting in Oregon in 1994, and previously observed patterns of recolonizing puma populations stabilizing after an initial growth period (Beausoleil et al. 2021; Oregon Department of Fish and Wildlife 2017; Ruth, Buotte, and Hornocker 2019). Additionally, though the population trend in recent years is less clear, pumas are still present on the landscape in significant numbers, with an estimate of 2.2 cougars 100 km^-2^ in Starkey during 2017 (Ruprecht et al., 2021).

### Process Model

The IPM is built around a process model which connects component models that estimate survival, recruitment, and abundance through a joint likelihood (Fig. 3b). The process model generates sex-specific abundance of calves and adults on November 1 of year *t* from abundance at year *t-1*, survival from year *t-*1 to *t*, recruitment of calves during year *t*, and net, human-mediated changes to the population between year *t-*1 and *t* (Fig. 3). We used November 1 to mark the biological year because it aligned with data collection, but it changes some traditional definitions of elk life history. For clarity, we refer to newborns as neonates until November 1 and calves during the following biological year. November was the first count of calves each year, so recruitment is the product of natality, e.g., the number of neonates born per female, and survival from birth to counting.

**Figure 3:**
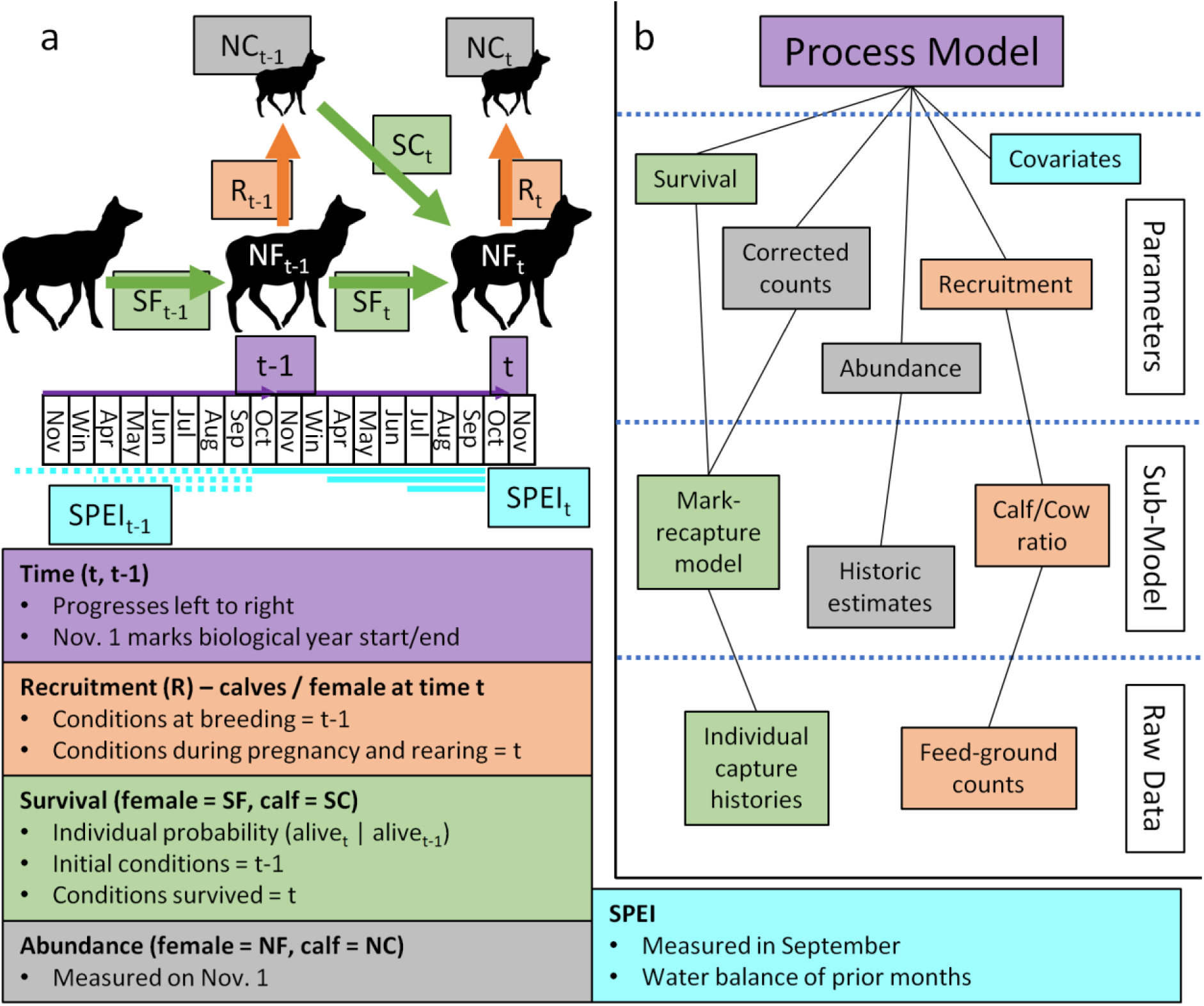
A conceptual diagrams of the IPM. (a) Timeline of the IPM process model over a single time-step. The boxes are color coded with purple representing time, orange representing recruitment, green representing survival, cyan representing covariates, and grey representing abundance. The small elk silhouette represents abundance in the calf stage (NC), and the large silhouette represents adult female abundance (NF). Recruitment (R – orange arrows) is the number of elk calves added to the population as a proportion of the adult females alive in that year. Calf survival (SC – slanted green arrow) is the probability of calves surviving from 0.5 years to 1.5 years old. Adult female survival (SF – horizontal green arrows) is the probability of adult females alive in November of year *t-*1 still being alive in year *t*. SPEI is used as an example covariate to clarify indexing. In this case, the cyan lines represent 3, 6, and 12-month SPEI, with the solid lines representing SPEI_t_ and the dashed lines representing SPEI_t-1_. (b) Diagram of the information flow in the IPM. The information flows from bottom to top, starting as raw data, being processed through a demographic sub-model into estimates for each parameter, which are then combined through the process model. The mark-recapture model is a Cormack-Jolly-Seber model used to estimate survival and detection probability. Detection probability is used to correct the count of the number of animals observed alive each year for imperfect detection, leading to the corrected counts that inform abundance. Cow/calf ratios are used to estimate recruitment in a binomial model. Recruitment, survival, and detection probability are estimated as independent variables each year in the sub-models. In the process model, survival and recruitment vary around a global mean rate. Covariates are not included until the combined process model.

The abundance of calves at year *t*, *N_calf,t_*, follows a binomial distribution with the total possible calves equaling the adult females that survived from *t-1* to *t*, *N_a,f,t-1_ × S_f,t_*, and recruitment per surviving female, *R_t_*:

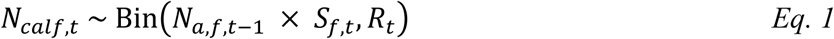

We assumed calves follow a 1:1 sex ratio based on previous elk calf capture and survival monitoring in the Blue Mountains of Oregon (Johnson et al., 2019; Kohlmann, 1999) and the years for which sex ratios of calves were recorded in Starkey. We defined adults as all non-calf individuals, which includes individuals less likely to reproduce (i.e., yearlings), because mature and yearling females are not distinguished in the population counts and historic abundance estimates. Adult abundance of sex *x* at time *t*, *N_a,x,t_*, includes four components: (1) the number of adults of that sex surviving from the previous year, (2) the number of adults of that sex harvested in the fall preceding November of year *t*, *N_h,x,t_*, and (3) the number of calves surviving from the previous year, and (4) the net change in each age/sex class due to management actions during the winter.

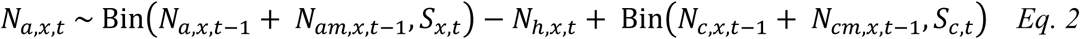

In equation 2, the first binomial term represents the surviving adults where *N_a,x,t-1_* is the abundance of adults of sex *x* in the previous November, *N_am,x,t-1_* is the net change adults due to management actions during the winter, and *S_x,t_* is the survival probability of adults of sex *x* from *t-1* to *t*. Similarly, the second binomial term is the number of calves surviving from *t-1* to *t* with a starting population of the calves recruited into the population in the previous year, *N_c,x,t-1_,* plus the net change in calves in the population from management actions in the previous year, *N_cm,x,t-1_*. The number of elk added or subtracted by harvest and management actions are known integers. In two years, 2017 and 2020, a substantial portion of the population was removed without fully recording the IDs. In all other years, the IDs of elk added or removed were fully recorded. To avoid biasing the demographic rates, we censored the survival estimates for 2017 and 2020 from the IPM (SI 2).

We used the normal approximation of the binomial distribution throughout the process model.

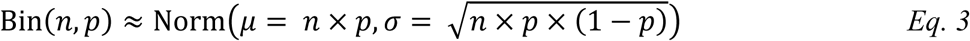

Using the normal approximation accommodates nonzero probabilities of obtaining negative values when subtracting the number of harvested individuals. To prevent negative populations, we truncated the normal approximations of the binomial at the minimum number of individuals known alive in each age and sex class based on feed ground attendance histories. The normal approximation of the binomial distribution is considered highly accurate when both the expected numbers of failures and successes are greater than or equal to 10 (Chang & Lee, 2017), which is met for every parameter in the IPM.

### Component Models

Component models that estimate survival, recruitment, abundance, and detection probability link the demographic factors in the process model to data in the joint likelihood of the IPM (Fig. 3b). The first component model is a Cormack-Jolly-Seber (CJS) model that estimates survival of males, females, and calves and detection probability of males and females (Cormack, 1972; Lebreton et al., 1992). CJS models condition on first capture, so calves do not have a detection probability. We built individual capture histories from observations at the feed grounds and harvest where *y_i,t_* = 1 if individual *i* is seen alive or harvested in year *t*, and *y_i,t_* = 0 if individual *i* is not seen. Detection is modeled as a Bernoulli random variable following

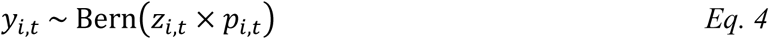

where *z_i,t_* is 1 if individual *i* is alive and 0 if individual *i* is dead, and *p_i,t_* is the probability of detecting individual *i* at time *t*, which is the same for all individuals of a sex within year *t* but varies with a fixed effect by year. The true state of individual *i* at time *t*, *z_i,t_*, is also a Bernoulli random variable following

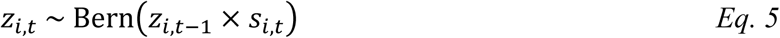

where *z_i,t-1_* is the state at the previous time step and *s_i,t_* is the probability that individual *i* survives from *t-1* to *t*, which is the same for all individuals of the same age and sex class within a year but varies by year. For each individual, *i*, we excluded values of *t* for which individual *i* was known to be unavailable (e.g. confirmed dead or documented as removed from the population as a management action).

We also use the CJS model to adjust the count of individuals encountered each winter for imperfect detection using

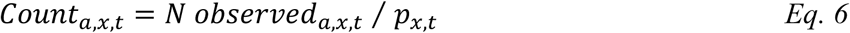

Here, *N observed_a,x,t_* is the number of adults of sex *x* observed on the feed grounds or through harvest in year *t*, and *p_x,t_* is the probability of detecting individuals of sex *x* in year *t* absent any individual variation, so *Count_a,x,t_* is a point estimate of the total number of adults of sex *x* in the population. This is similar to a Jolly-Seber estimate of abundance, however, because we do not necessarily meet the equal capture probability between seen and unseen individuals, we do not include information on the precision of the estimate (Jolly, 1965; Seber, 1965). Instead, we include the count as data in the IPM with a normal error following

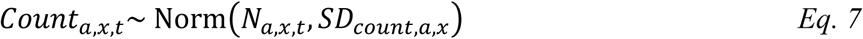

where *SD_count,a,x_* quantifies the difference between the counts and the IPM-estimated abundance, *N_a,x,t_* (Schaub & Kéry, 2021). We used a diffuse normal prior truncated to positive values for each age-sex class’s standard deviation of counts. The values for each count standard deviation are estimated only from the joint likelihoods in the IPM; no information about the precision of the counts was passed to the IPM. Count data included without precision is the backbone of many basic IPMs (Schaub & Kéry, 2021). Dividing the naïve counts by the detection probability serves only to mitigate the bias of imperfect detection.

In addition to its inclusion in the process model, recruitment in year *t*, *R_t_*, is estimated using the count of calves on the winter feed grounds, *Count_calf,t_*, as a binomial random variable with the number of draws equal to the count of adult females, *Count_a,f,t_*, and probability *R_t_*.

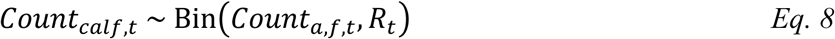

Lastly, we incorporated the existing population estimates from sightability-corrected aerial surveys and mark recapture surveys (Rowland et al., 1997; Unsworth et al., 1990, ODFW, unpublished data) into the IPM as data. In this formulation, the mean of the estimate, *N-est_a,x,t_*, is incorporated as a normal random variable with mean of *N_a,x,t_,* which is generated by the process model, and with the standard error estimated from the aerial surveys, *SE-est_a,x,t_*.

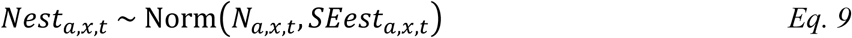

In 2 years, 1996 and 2005, both sightability and mark recapture estimates are available for at least one age-sex class. In those years, we passed both as independent estimates.

### Covariate Modelling

We modeled two demographic rates, *R_t_* and *SC_t_*, as a function of the covariates predicted to correlate with elk demography based on our hypotheses (*Table 1*) using a logit link function in the combined, process model. Using *R_t_* with the climate covariate *SPEI* as an example, the covariate effects follow

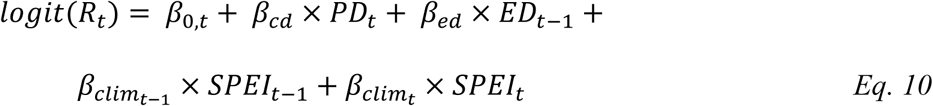

*B_0,t_* is the intercept for year *t*, *PD* is puma density, *ED* is elk density, and *SPEI* is the precipitation/evapotranspiration index (though we tested versions with each of the density-independent, bottom-up covariates in Table 3). We used both SPEI from the current year, *SPEI_t_*, and previous year, *SPEI_t-1_*, as model inputs (Fig. 3a, Table 2). For *R_t_*, *SPEI*_t-1_ represents drought reducing maternal body condition, leading to lower conception rates, *SPEI_t_* represents nutritionally stressed neonates dying before calves are counted in the fall, *ED_t-1_* represents competition for forage limiting female body condition prior to breeding thus reducing pregnancy rates, and *PD_t_* represents predation on neonates prior to fall counts. For *SC_t_*, *PD_t_* represents predation between 0.5 and 1.5 years, *ED_t-1_* represents decreased survival through nutritional limitation of nursing neonates before entering winter due to competition for forage, *SPEI_t-1_* represents poor nutrition entering winter, and *SPEI*_t_ represents poor over-summer nutrition. We implemented the covariate effects in the process model which combines all of the demographic estimates (Fig. 3b). In the component models, we estimated each demographic rate independently by year. In total, we fit 9 versions of the IPM: one with each of the climate covariates paired with the mean of the puma density estimates, the top-performing climate covariate paired with the logistic reconstruction of puma density, and a null version with no covariates. The covariate runs all used the median elk density from the null model for *ED_t-1_*.

### IPM Implementation

We implemented the IPM in two stages. We first ran each of the component models separately and then used the means and standard deviations from the posterior distribution of each parameter to define the distribution of possible values for that parameter (Moeller et al., 2021). This method of incorporating information from the component models provides two benefits. First, it is much more computationally efficient, allowing the IPM to be run in hours rather than weeks. Second, it is the same method used to include the historic abundance estimates, meaning all of the information in the IPM is incorporated using a consistent approach (Schaub & Kéry, 2021). Tests on smaller versions of the data revealed no differences in the estimates from this approach and a fully integrated approach for the first 24 years (SI 2). In the final 12 years, we violated the closure assumption of the CJS model by removing a large proportion of the population without recording IDs. To avoid biasing all of the survival estimates through the random effects structure, we censored those years in the two-stage version of the IPM. We did not censor those years in the fully integrated tests, because doing so would have required additional parameters and affected precision throughout the model. The differences in the underlying data lead to predictable differences in the survival estimates (SI 2).

In the component models, we estimated each parameter independently for each year such that *R_t_* can take any value from 0 to 1 for all *t,* and annual values are not linked to each other. We used priors that are flat on the probability scale, either Beta (1,1) or Logistic (0,1), depending on whether the model was written to require a transformation from the real scale, for all parameters (Northrup & Gerber, 2018). In the component models, we estimated independent annual values for recruitment, calf survival, female survival, male survival, and detection probability of males and females.

The second stage of implementing the IPM is running the process model with the results of the component models and historic estimates incorporated as data (Fig. 3b). In this stage, demographic parameters are treated as varying annually around some mean value following

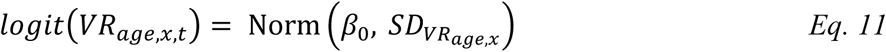

where *VR_age,x,t_* is a vital rate of age class *age* and sex *x* in year *t* (*e.g.* adult female survival in 1990), *β_0_* is the mean of that vital rate on the logit scale, and *SD*_*VRage*,*x*_ is the standard deviation of the annual values of the vital rate. Adult survival was the only parameter with an informative prior, *Beta* (3, 1.2), which we chose qualitatively as a prior that gave more weight to high survival probabilities without making low survival probabilities impossible to stabilize adult survival estimates in years with minimal or no data. We implemented the component models using NIMBLE in R (NIMBLE Development Team, 2024; R Core Team, 2024) and the IPM in JAGS (Plummer, 2003) (see SI 2 for the reasons behind the sampler choices). We ran three chains of the IPM for 1,000,000 iterations after an adaptation phase of 10,000 iterations and a burn-in of 100,000 iterations. We thinned the results by 100. The component models converge and mix much more quickly. We ran three chains of each component model for 25,000 iterations after a burn in of 10,000 and did not thin the posteriors.

### Population Growth

To test whether the hypothesized drivers of elk demography affect population growth, we estimated population growth as a function of survival and recruitment. We could not calculate lambda as *N_t_* / *N_t-1_* because harvest and management additions and removals would mask the natural variation. Instead, we constructed a female-only matrix model (Leslie, 1945) using the values for recruitment, calf survival, and female survival each year at each step of the MCMC. We then calculated what lambda would be without human action as the dominant eigenvalue of the matrix. To assess the influence of the covariates on population growth, we constructed the same matrix using the global means of recruitment, calf survival, and female survival at each step of the MCMC. We then varied one covariate while holding all others at 0 (the centered and scaled mean), obtaining a posterior distribution of population growth under the full range of values for each covariate. We defined the effect of the covariate as the slope of population growth for a standard-deviation change in the covariate.

### Pregnancy Rates

To elucidate mechanisms influencing recruitment, we separately modeled pregnancy probability as a function of female density, *ED_t_*, and 12-month SPEI from the summer immediately preceding breeding, *SPEI_t_*. To limit our hypothesis tests, we did not run models using any climate covariates other than 12-month *SPEI_t_*, which performed the best in the IPM (see results and SI 1). We also did not test for lag effects, e.g. *SPEI*_t-1_, because we did not hypothesize a mechanism for carryover effects on pregnancy probability. Lactating females, young females (< 3 years-old), and old females (> 13 years old) are all less likely to get pregnant than prime-aged, non-lactating elk (Piasecke et al., 2009), so we modelled each age × lactation class separately following the form:

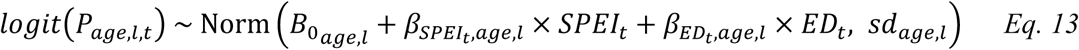

where *P_age,l,t_* is the probability of an individual in age class *age*, and lactation status *l*, being pregnant in year *t*. We estimated *P_age,l,t_* in a binomial model where success were the number of pregnant females and trails were the number of females tested for pregnancy status. We implemented the pregnancy model in NIMBLE and R (NIMBLE Development Team, 2024; R Core Team, 2024) using the same run parameters as the IPM component models.

## RESULTS

The IPM converged well and traced expected trends in the elk population including a declining population during heavy harvest of adult females from 1998 to 2007, an increasing population after adult female harvest ceased, and declines again following the two intentional elk population reductions in 2017 and 2020 (Fig 4). Mean adult female survival probability was 0.93 (95% credible interval: 0.91 – 0.95), mean calf survival (0.5 to 1.5 years) probability was 0.79 (0.70 – 0.88), and mean adult male survival was 0.81 (0.75 – 0.88). Survival terms all exclude harvest mortality. Mean recruitment was 0.36 (0.32 – 0.40) calves per adult female. The geometric mean of population growth was 1.03 (1.01 – 1.04). For females between 3 and 13 years-old (prime-aged), mean probability of being pregnant was 0.92 (0.9 – 0.95) when not lactating and 0.88 (0.82 – 0.92) when lactating. For females > 13 years old, the annual mean pregnancy probability was lower – 0.75 (0.64 – 0.85) when not lactating and 0.70 (0.57 – 0.83) when lactating. Adult females < 3 years-old, all of which were non-lactating, had an annual mean pregnancy rate of 0.94 (0.87 – 0.98), which is likely skewed high due to 15 years of low sample sizes (≤5) and pregnancy rates of 1 (naïve pregnancy rate among non-lactating, young female when pooling all years was 0.88, SI 3). Lactating young females were excluded due to low sample sizes (zero in most years, see Methods). To report the evidence for a nonzero effect of each covariate on demographic quantities within the Bayesian models, we used the posterior probability, P(d), that the covariate correlated with the parameter of interest in the direction predicted by the corresponding hypothesis. Thus, a P(d) near one indicates support for a hypothesis, a P(d) near 0.5 indicates no correlation, and a P(d) near 0 indicates correlation in the opposite direction predicted by the hypothesis.

**Figure 4:**
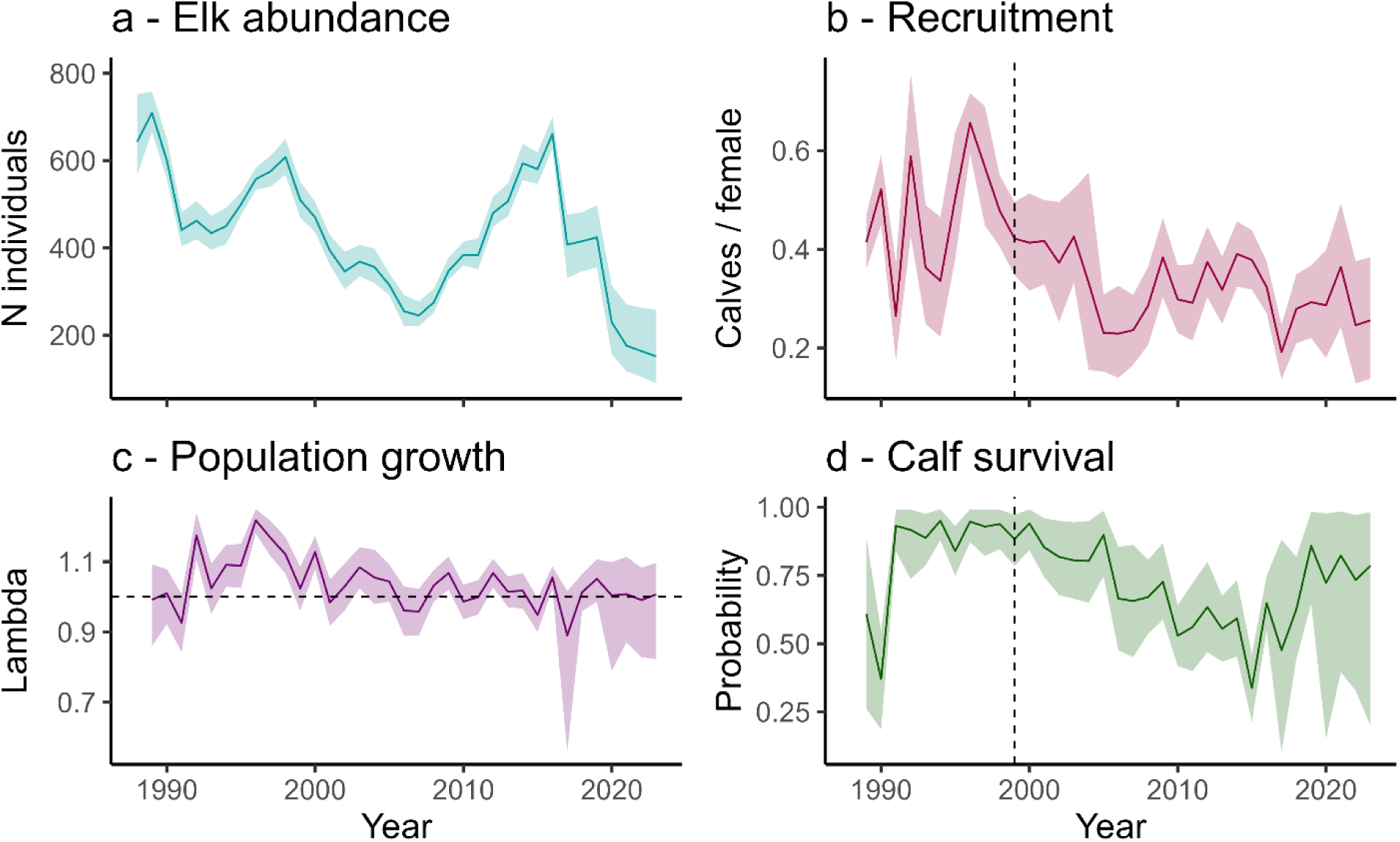
The trends in (a) abundance including all age/sex classes, (b) recruitment, (c) population growth, and (d) calf survival (6 months to 18 months) through time. The solid lines represent the median of the posterior distributions, and the colored bands represent 95% credible intervals. In the recruitment and calf survival plots (b and d), the vertical dashed line marks 1999, the year that the logistic growth model of puma density reached half carrying capacity. In the population growth plot (c), the horizontal dashed line marks a value of one, indicating a stable population.

We found evidence of bottom-up limitation through climate in some, but not all, of the covariates representing density-independent, bottom-up effects. Of the lag effects tested for recruitment, e.g., *Cov_t-1_*, we found positive effects of *PDSI_t-1_* and 12-month *SPEI_t-1_* on recruitment (P(d) of 0.97 and 0.99 respectively) and a negative effect of mean summer *temperature_t-1_* (P(d) = 0.98). We also found weak evidence for an effect of 6-month *SPEI_t-1_* (P(d) = 0.94). Of the same-year effects tested for recruitment, e.g., *Cov_t_*, we found a negative effect of *NDVI_t_* (P(d) of 0.02) and weak evidence for a negative effect of *precipitation_t_* (P(d) = 0.10) and a positive effect of 3-month *SPEI_t_* (P(d) = 0.91). The negative effects of *NDVI_t_* and *precipitation_t_* are the opposite of the predicted direction under the density-independent, bottom-up hypothesis. None of the other climate covariates showed an effect on recruitment. For climatic effects on calf survival, we found weak evidence of a positive effect of 12-month *SPEI_t-1_* (P(d) = 0.91) and of negative effects of *temperature_t_* and *temperature_t-1_* (P(d) of 0.07 and 0.06 respectively). The negative effects of temperature are the opposite of the predicted direction, while the positive effect of 12-month *SPEI_t-1_* supports the hypothesis that summer drought while a neonate reduces overwinter survival of calves. No covariates showed an effect on calf survival at an alpha of 0.05. Of the bottom-up covariates, 12-month *SPEI_t-1_* had the largest effect and the highest probability of a directional relationship with both recruitment and calf survival. The posterior effects of puma and elk density were similar across models using all climate covariates (SI 1), so, for brevity, we focus on the model using 12-month *SPEI* throughout the rest of the results. The coefficient on recruitment for 12-month *SPEI_t-1_* had a posterior median of 0.22 (0.05 – 0.39), indicating that wet conditions prior to the breeding season were associated with increased recruitment (Fig. 5a3). Specifically, a 1-standard deviation increase in 12-month *SPEI_t-1_* correlated with a 24% (5 – 48%) increase in the odds of female elk successfully recruiting a calf in November. It also had a positive correlation with pregnancy rates in lactating, prime-aged (0.62; 0.01 – 1.21; P(d) = 0.98) and old elk (0.51; −0.21 – 1.29; P(d) = 0.92), increasing the odds of pregnancy in those groups by 86% (10 – 235%) and 67% (−20% – 263%) respectively. The weak evidence of a positive relationship between 12-month *SPEI_t-1_* and calf survival had a posterior median of 0.36 (−0.16 – 0.96; P(d) = 0.91), corresponding to a 43% (−15% – 161%) increase in the odds of survival per standard deviation change in 12-month *SPEI_t-1_*. The effect of 12-month *SPEI_t-1_* on elk vital rates translated to a positive effect on population growth, with a slope estimate of 0.02 (0.01 – 0.04; P(d) > 0.99) on population growth. In total, these results show drought acting in two ways to decrease elk population growth: first by lowering female body condition during breeding, thus reducing pregnancy rates, and second by limiting neonate growth prior to winter, thus reducing calf survival.

**Figure 5:**
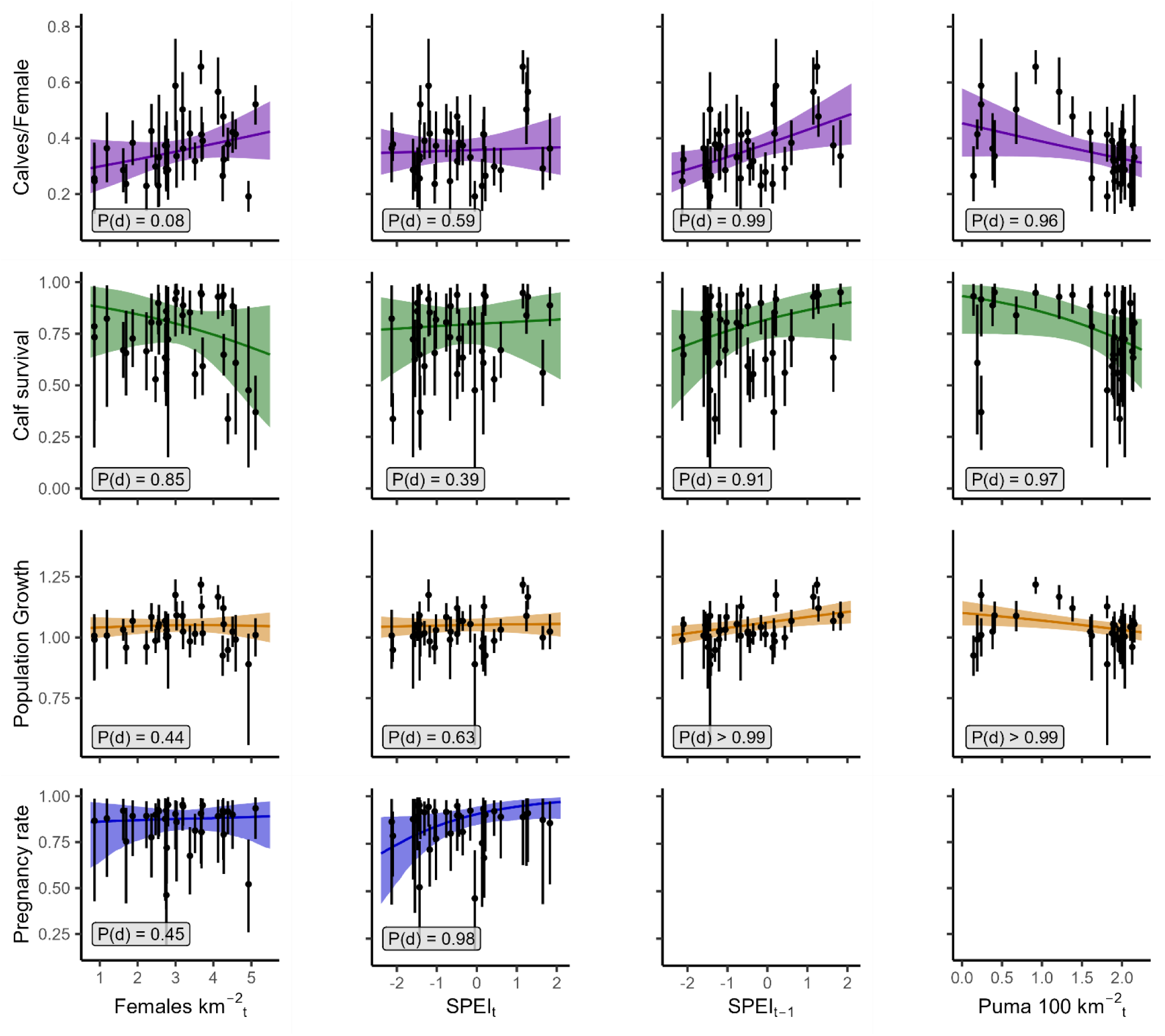
The marginal plots for the variables on recruitment (first row) and survival (second row) in the IPM, population productivity, *PP* (third row), and pregnancy probability of lactating females between 3 and 13 years-old (fourth row, see SI Fig 3.6 for the other pregnancy marginal plots). The first column is female density, representing density dependence. Density dependence was measured at the time of mating in all models. The index *t* is correct for the x-axis of pregnancy model that it appears on, but would be *t-1* for all other models. The second and third columns are the Standardized Precipitation and Evapotranspiration Index from the current and previous year (SPEI_t_ and SPEI_t-1_) respectively, representing forage conditions. The fourth column is puma density, representing predation pressure. The final two plots are blank because we did not hypothesize mechanisms for the either the previous year’s drought index or predation affecting pregnancy probability of lactating females. The numbers in each graph are the probability of a correlation in the predicted direction.

We also found evidence for top-down forces on elk demography. The scaled mean of the two puma density estimates was negatively related to elk recruitment with a coefficient of −0.19 (−0.40 – 0.02; P(d) = 0.96; Fig 5), which represents 17% decrease in the odds of recruitment per standard deviation increase in puma density. Puma density was also negatively correlated with calf survival from 0.5 to 1.5 years with a coefficient of −0.60 (−1.35 – 0.03; P(d) = 0.97; Fig 5), or a 45% decrease in the odds of survival. Those effects combined to cause a 0.03 (0.04 – 0.01 P(d) < 0.99) decrease in population growth per standard deviation increase in puma density. Using the logistic reconstruction of puma density, we estimated slightly larger effects of puma density for both recruitment and calf survival (SI 1). The coefficient on recruitment was −0.23 (−0.40 – −0.05; P(d) = 0.98), which corresponds to a 20% decrease in the odds of recruitment per standard deviation increase in puma density. The coefficient on calf survival was −0.67 (−1.28 – −0.14; P(d) = 0.98), which corresponds to a 49% decrease in the odds of survival from 0.5 to 1.5 years. Both versions of the covariate indicate that puma density acts to reduce calf survival during both the neonate and calf stages.

We found no evidence for negative effects of density dependence. Adult female elk density the previous year did not affect calf survival (P(d) = 0.15; Fig. 5b1) in the IPM, and may have had a positive effect on recruitment, with a beta estimate of 0.13 (−0.06 – 0.32; P(d) = 0.92). We also did not detect an effect of density dependence on population growth or pregnancy probabilities. The density dependence effect on population growth was 0.002 (−0.02 – 0.02). The pregnancy probability of non-lactating, old females may negatively correlate with female density (−0.36; −0.94 – 0.16; P(d) = 0.91), but no other age/lactation class had a P(d) > 0.8.

## DISCUSSION

Using 36 years of data, we identified both bottom-up and top-down influences on elk demographic rates that had additive effects on elk population growth. Quantifying those effects required a study that spanned the recovery of a major carnivore, 5-fold changes in elk density, and three and a half decades of weather variation characterized by intensifying drought. Our results highlight that, without long-term studies, we cannot fully explore the forces acting on population dynamics. That is particularly true in large herbivores for which in-situ experiments are cost prohibitive or untenable, leaving only observational studies. Study length directly determines which mechanisms are testable in single-site, observational studies by limiting which parameters undergo variations large enough to measure demographic responses. Study length also limits the statistical power of population-scale demographic studies. Each year yields a single data point for a demographic rate, meaning that short-term studies can identify only the largest effects and do not have the power to test multiple hypotheses simultaneously. Exploiting spatial variation can mitigate those limitations, but studying processes across broad geographic ranges also introduces the potential for confounding, between-site variation that must be identified and modelled. Space-for-time substitutions also fail to identify trends within sites, such as the drought and predation driven decline in recruitment and calf survival we measured.

The declines we measured in recruitment and calf survival after puma recolonization are both plausibly explained by consumptive effects of puma alone. The 27% decline in elk recruitment after puma recolonization indicates that, at the average female abundance (228 individuals), 27 fewer neonates survive the 22 weeks between a June 1 birth and a November 1 count. Ruprecht et al. (2021) estimated puma density in Starkey in 2017 to be 2.2 adult cougars 100 km^-2^. At that density, a kill rate of ∼0.7 elk neonates per puma per week would fully account for the decline in recruitment after puma recolonization. Estimated kill rates for pumas preying on deer and elk during summer range from 1 to 1.75 kills per week (Cooley et al., 2008; Ruth et al., 2019), putting the decline in survival to 6 months well within the range that can be explained by consumptive effects. In Starkey, elk outnumbered deer for the entirety of our study (Rowland et al., 1997; ODFW and USFS, unpublished data), and 61% of puma scats collected in late spring/early summer contained elk while less than 20% contained deer (data from 2017; Ruprecht et al. 2021). In Eastern Oregon, puma selected for elk neonates from late May through October even in a system with more deer than elk (Clark et al., 2014). The magnitude of the change in calf survival following puma recolonization is similarly plausible based on consumptive effects alone. At the mean estimated calf abundance (110 individuals), a kill rate of 0.29 elk between 0.5 to 1.5 years-old per puma per week would explain the decrease in calf survival. These results also match the literature, which has found limiting effects of puma abundance on elk calf survival and recruitment (Johnson et al., 2019; Proffitt et al., 2014, 2020).

In addition to puma, both wolves and black bears have increased in the area since the start of the study, but their effect on elk demography was likely limited compared to pumas. The Ukiah wolf pack was not established in Starkey until 2021 and so could not have influenced the inference from this time series. From 2018 onward, non-territorial wolves were infrequent visitors based on detections on the Starkey camera grid, suggesting limited impact on elk during our study (ODFW unpublished data, Ruprecht et al. (2021) describes the camera grid). Black bears were likely never extirpated from the Starkey area and did increase in abundance through the course of our study (ODFW, 2012). However, black bear predation of elk in the first 6 months of life is limited compared to pumas in the Blue Mountains and falls off dramatically after the first few weeks post-birth once calves are mobile enough to stay with their mothers (Johnson et al., 2019). Indeed, the total effect of black bear predation on elk calves across Oregon is largely compensatory while puma predation is at least partially additive (Johnson et al., 2019). Further, black bears should not affect calf survival in our study, which we measure from 6 months to 1.5 years, because calves older than 6 months are relatively invulnerable to black bear predation. Finally, a meta-analysis of studies across the western U.S. found that puma outpace black bears as a source of cause-specific mortality in elk over the first 3 months post-birth in systems without wolves or grizzly bears (though black bears lead in the first month) (Griffin et al., 2011).

Predation could also explain the weak evidence we found for positive density dependence in recruitment. Invariant predation pressure can lead to positive density dependence as prey populations fluctuate. At high prey density, this occurs through predator swamping (Darling, 1938; Ims, 1990). Under predator swamping, the number of elk neonates that predators can consume in a summer has a maximum threshold. Neonates born in excess of that threshold survive to November at higher rates, leading to more calves per female at greater elk abundances if assuming stable fecundity. At low prey density, positive density dependence occurs if predators do not numerically respond to the decline in prey density and per-predator consumption remains high (Holling, 1959; Sinclair & Pech, 1996). In that scenario, the total number of neonates consumed is constant, but the proportion consumed increases as the number born decreases. In a recent elk reduction in Starkey, neither puma nor black bear showed evidence of a numeric response (Loonam et al., in prep). If these species did not prey switch, the lack of numeric response could lead to positive density dependence in recruitment.

Summer precipitation can have important implications for elk nutritional condition and pregnancy rates in ecosystems characterized by dry summers (J. G. Cook et al., 2004, 2018; Johnson et al., 2013; Lukacs et al., 2018) but tends to have minimal effects on elk survival (Brodie et al., 2013; Johnson et al., 2019). Our study supports these patterns around pregnancy rates, with both pregnancy and recruitment declining in drought conditions. With an average of just under half of the females lactating each year, the magnitude of the decline in pregnancy rates of lactating females during drought can fully explain the decline in recruitment. However, we also found weak evidence of calf survival declining with drought (Fig. 5). Specifically, survival from 0.5 to 1.5 years-old was lower when the preceding summer, i.e., the summer spent as a neonate, was characterized by more drought-like conditions. This outcome is likely driven by slower neonate growth in summers with poor forage, leading to smaller calves which have lower over-winter survival probabilities (J. G. Cook et al., 2004). With drought already affecting elk population growth through both pregnancy rates and calf survival, the total effect of drought on elk population growth warrants further investigation.

As climate change continues, drought conditions will become more frequent and severe in the western U.S., likely expanding past the climatic variability observed in our study period and shifting vegetative communities to less nutritious compositions (Dalton et al., 2021; Halofsky et al., 2018; Hicke et al., 2022; Kerns et al., 2018). In our study system, the trend in increasing drought is already measurable, including a significant trend over the course of our study alone (Fig. 2). This change explained a 3% decline in elk population growth over 36 years. That decline was likely caused by measurable changes to growing season conditions in our system, including shortening of the growing season by 2 weeks and plant senescence advancing by 2-3 weeks between the 1990’s and 2020’s (Brown et al., 2022). This decline in forage availability and compressed growing season will likely continue with climate change (Dalton et al., 2021) and may increase the importance of weather patterns on elk demography. Elk body condition (Brown et al., in prep), pregnancy of lactating females, and calf survival already show measurable effects of drought. Further work should identify the thresholds at which adult survival and pregnancy rates of non-lactating females begin to decline with drought. Responses in those vital rates would significantly increase the magnitude of drought effects on population growth.

While both predator recovery and climate change shaped variation in elk demography over the past 36 years, climate change is likely to dominate annual variations going forward. Once recovered, populations of large carnivores tend to stabilize. At that point, the additive effects of predation will persist but act at relatively constant levels from year to year. In contrast, conditions driven by climate change, such as drought in the western U.S., will continue becoming more common and more extreme in the coming years. In our study, the acceleration of drought over 36 years nearly matched the total effect of the recovery of a major predator on elk population growth. Even if the effects of climate change on elk demography continue linearly, that rate of decline from bottom-up forces could outpace the effect of an intact, North American carnivore guild within decades (though the continued expansion of wolves and the shelter from predation that elk get in the feed grounds in winter could mean that predation has larger effects than estimated here). If the more stable, but more influential, demographic rates, such as adult female survival, begin to decline with more severe climate change, the shifts could be even more rapid. This pattern is likely occurring in flagship herbivores across the globe but being masked by compensatory life history strategies and the paucity of long-term studies. Nutritional limitation in large herbivores acts on body condition and pregnancy probabilities first. Those effects can be mitigated through every-other-year reproduction until the nutritional limitation is severe enough to impact adult survival or pregnancy rates of non-lactating females. With the high nutritional needs and slow life histories of large herbivores, recovery from even temporary lean times may be difficult as they become more frequent.

## Supporting information

Appendix S1 - Covariates

Appendix S2 - IPM Formulation

Appendix S3 - Residual Plots

## ACKNOWLEDGMENTS

Funding for this research was provided by the U. S. Forest Service, Pacific Northwest Research Station, the Oregon Department of Fish and Wildlife, and Federal Aid in Wildlife Restoration grants. Many ODFW and USFS employees made this publication possible through countless hours of data collection, project management, animal handling, and innumerable other activities that have kept the Starkey Project running since 1988. Of particular note for this paper are H. Martin, T. Forrester, C. Eckrich, M. Bianco, B. Naylor, J. Smith, N. Libal, C. Borum, B. Dick, D. Rea, and R. Kennedy. B. Johnson provided friendly reviews throughout the process and shared invaluable expertise from his decades working in Starkey and similar systems. J. Ruprecht was an excellent friend and sounding board throughout the process and provided insightful reviews of the manuscript. The coding and IPM implementation would have taken much longer without the help of A. Moeller getting us started. Levi was supported by NSF Award 2317537.

