## Appendix S1 - Covariates for "The effects of predator recovery and climate change on the long-term demography of a flagship herbivore"

### SI 1 – Covariate Selection and Comparison

*1.1 – Puma Covariate*

Figure S1.1: Two data sources for puma density around Starkey and two methods of combining the sources. All parameters were centered and scaled using the range of years for which they are all available and show the same trend (1994 through 2012). The purple crosses are a population reconstruction from the Starkey Wildlife Management Unit, the finest scale for which puma density estimates were available. The population reconstruction will be biased low in recent years because puma deaths have not filled in the population in those years yet. The blue squares represent ODFW's estimate of puma abundance in the Blue Mountains Region that contains the Starkey Wildlife Management Unit. The green triangles represent a smooth logistic growth of the population fit to the population reconstruction data from 1987 to 2012. The red circles are the mean of the ODFW estimate for the Blue Mountains and the population reconstruction for the Starkey Wildlife management unit. The mean is used in the analyses in the main text.

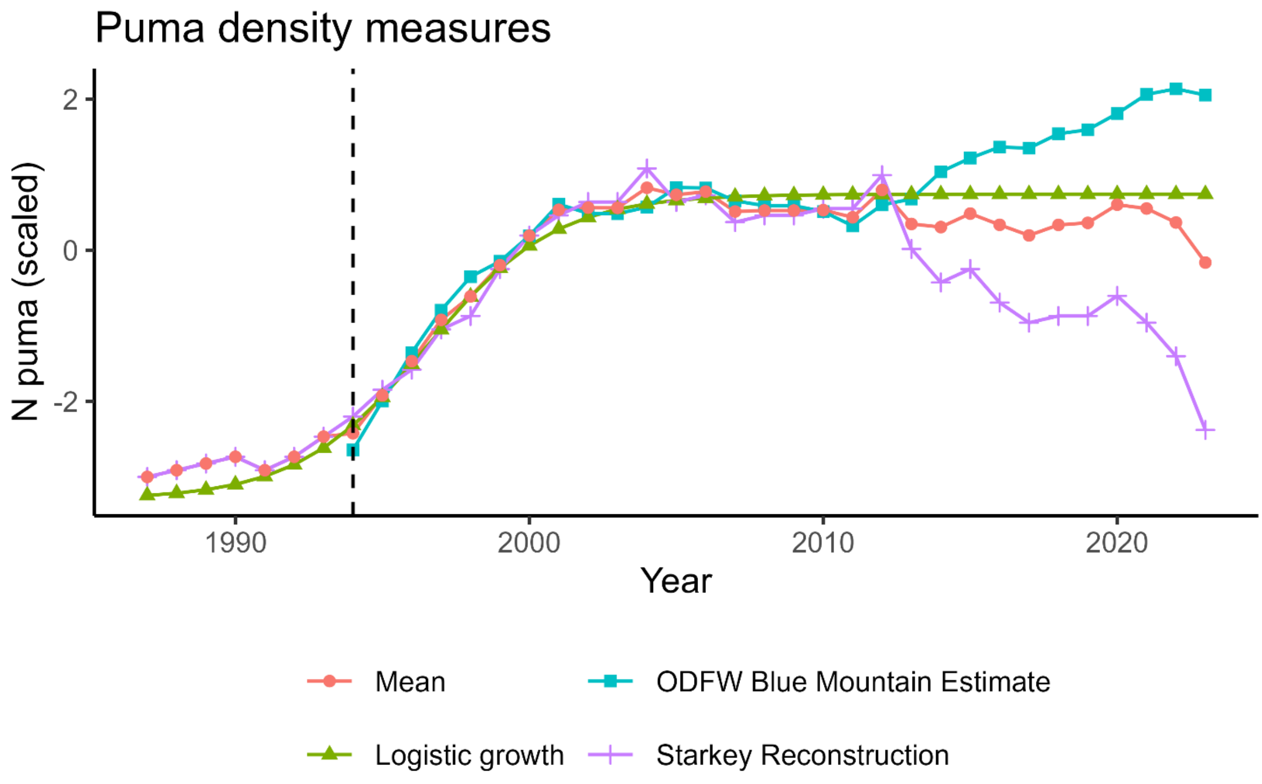

No estimate of puma abundance that spans the study timeframe exists for the study area, however ODFW estimates puma density using two methods at two spatial scales that encompass Starkey (Fig. S1.1). The first method is a simple population reconstruction for the Starkey Wildlife Management Unit using all known cougar mortalities (Clark, unpublished data). In the reconstruction, puma are aged upon death and added to the population for each year that they were alive. Because puma must die to be counted, the population estimate will be biased low in recent years. That lag time will be somewhere between 5 and 10 years, the 75$^{th}$ and 95$^{th}$ percentiles for age at death respectively. The second method is ODFW's estimate for puma density in the Blue Mountains (which encompass the Starkey Wildlife Management Unit). The ODFW estimate utilizes mortality and hunter success data in conjunction with a puma population process model to estimate puma density (Keister & Van Dyke, 2002). The two data sources show the same general trend in the population from 1994 (the year the ODFW estimate begins) through 2012, including the rate of puma increase and the timing of the population levelling out, however they diverge after 2012, with the reconstruction showing a decline and the estimate showing renewed growth.

With two estimates showing contradicting trends from 2013 through 2023, we worked to combine them for the final covariate. In the first version of the manuscript, we fit a simple logistic curve to the reconstruction data from 1987 through 2012 and extrapolated it out through the end of the study. That approach assumed that puma density levelled off and was steady after recovery, a pattern supported by the literature (Beausoleil et al., 2021; Oregon Department of Fish and Game, 2017; Ruth et al., 2019) and by the consistency of reported puma mortalities in the Starkey Wildlife Management Unit and the Blue Mountains (ODFW unpublished data). In this version, at the suggestion of a reviewer, we scale the two puma estimates using the mean and standard deviations from the years during which they show the same pattern, placing the two estimates on the same scale, then take the mean of the two scaled estimates, resulting in the line labeled mean in Fig. S1.1. The previous version of the manuscript found negative relationships between puma density and elk recruitment, calf survival, and population growth, regardless of the bottom-up covariate tested. Additionally, a version of the IPM from the early testing a troubleshooting stages found a negative relationship between the unedited population reconstruction and elk recruitment in a univariate version of the IPM. At that time, calf survival had no covariates tested and the follow-up regression on population growth was not conducted. We report those previous versions of the covariate for transparency but focus exclusively on the mean of the two estimates in the main text to avoid testing multiple combinations of covariates. We also found minimal differences in the effect estimates in the model using 12-month SPEI (Fig. S2.1)

Figure S2.1

Estimates of the beta values for covariates on recruitment (top panel) and calf survival (bottom panel) using the mean of the puma density estimates (left groups) and the logistic growth curve (right groups) as the puma covariate in the 12-month SPEI model. The model using the mean of the puma density estimates is the model focused on in the main text. When using the logistic growth curve, puma density has a larger estimated effect and SPEI has a smaller estimated effect.

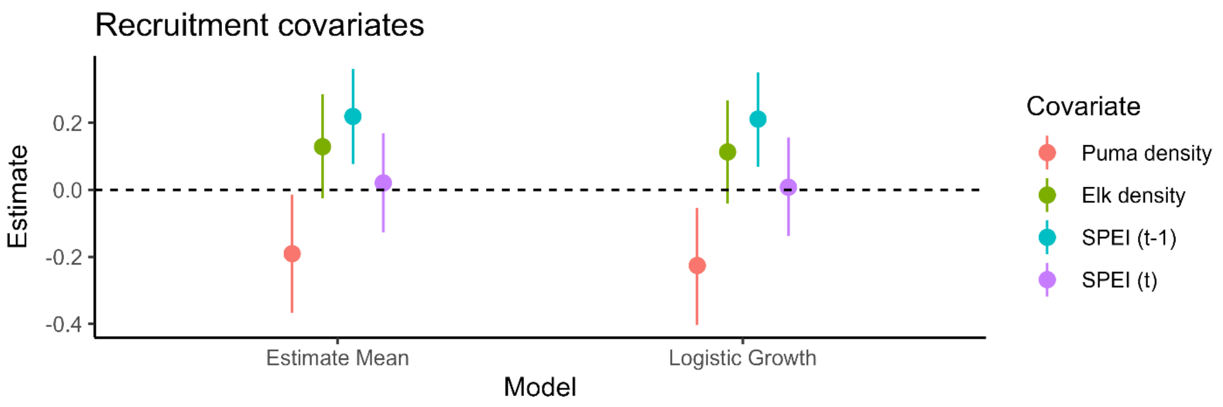


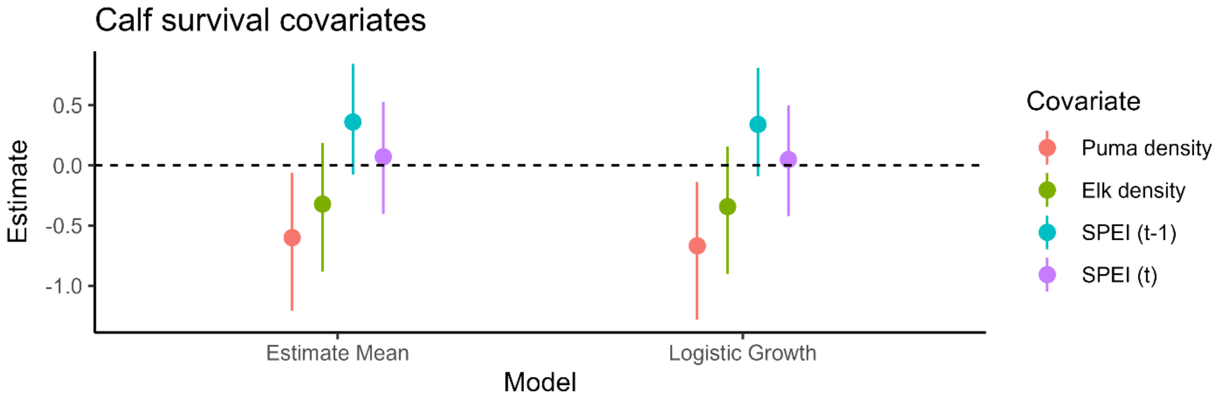


*SI 1.2 – Climate Covariates*

In all, we examined 7 covariates for potential density-independent, bottom-up effects: mean growing season temperature, total growing season precipitation, mean growing season normalized difference vegetative index (NDVI), mean growing season Palmer Drought Severity Index (PDSI), and the Standardized Precipitation and Evapotranspiration Index (SPEI) in September at three time-scales (3, 6, and 12 months).

For temperature and precipitation, we used Parameter-elevation Regressions on Independent Slopes Model (PRISM) data accessed from the “explore” tool on https://prism.oregonstate.edu on September 15, 2024 (PRISM Climate Group, n.d.). We downloaded the data for the monthly mean temperature and total precipitation for the centroid of Starkey from January 1, 1987 through January 1, 2024. We subset the data of each year to May through September, inclusive, and took the mean of the temperature and the sum of the precipitation for the final covariates.

For NDVI, no source provides reliable estimates for the entirety of our study, so we combined data from two sources: the Advanced Very High Resolution Radiometer sensors (AVHRR – GIMMS NDVI 3rd generation) and the Moderate Resolution Imaging Spectroradiometer Terra Satellite (MODIS – MOD13A2 V6.1 NDVI) (NOAA, NASA Earth Data). For both sources, we used the 16-day composite values and accessed the data through Google Earth Engine on September 15, 2024 (Gorelick et al., 2017). We used MODIS values for all years that it is available (2000 on) and transformed AVHRR values to the MODIS scale using a simple linear transformation based on values from a regression between MODIS and AVHRR values in our study area from 2000 to 2013, when AVHRR loses reliability ($R^2 = 0.875$). We could not consider any trimming to account for canopy cover due to the low resolution of AVHRR data. However, only ~10% of Starkey is closed canopy (ODFW, unpublished data), and canopy cover accounts for relatively little of the temporal variation in NDVI in landscapes with evergreen canopy (Gamon et al., 1995).

For PDSI, we acquired data from the National Centers for Environmental Information (NCEI NOAA; https://ncei.noaa.gov/access/monitoring/historic-palmers) on September 15, 2024. At the time of writing, the website is unavailable due to Hurricane Helene. In the first version of the manuscript, we used the PDSI value in September of each year as the covariate. In this version, we use the mean PDSI from May through September per reviewer suggestion. PDSI is cumulative, so PDSI in September incorporates information from the prior months (Palmer, 1965), resulting in the two measures being highly correlated ($R^2 = 0.761$) with the growing season mean putting more weight on conditions earlier in the season.

We acquired SPEI using the global drought monitor tool at https://spei.csic.es/ on September 15, 2024. SPEI measures the deviation of the precipitation/evapotranspiration balance over a given time interval from the mean precipitation/evapotranspiration balance over a reference period, in this case 1950 to 2010 (Vicente-Serrano et al., 2010). For time intervals, we chose the 3, 6, and 12 months preceding September of the year in question. We chose those 3 timeframes a priori to minimize the probability of false positives and to limit our investigation to the proximate effects of annual weather, rather accumulated, long-term effects.

*SI 1.3 – Models run*

We ran 8 models for inference in our investigation, a null model and a model using each of the 7 density-independent climate covariates (additional models that vary the model structure, rather than the covariates used, are detailed in SI 2). Each model with covariates included the puma density covariate described in SI 1.1, an estimate of elk density derived from the null model, and a climate covariate from both time *t* and *t-1*, representing the two time scales weather could act on to influence elk demography. The same covariates were used for both recruitment and calf survival. We found no difference between the demographic estimates among the models (Fig. S1.2) and little variation in the non-climate covariate estimates (Fig. S1.3). Of the climate covariates, 12-month SPEI had the largest estimated effect (Fig. S1.3) and is the model we focus on in the main text.

Figure S1.2

Selected vital rates through time from each of the 8 models. The 8 models are defined by the climate covariate used. In the years that the estimates of recruitment and calf survival diverge, each estimate still falls within the middle 50% of the posterior of each of the other models. Plotting the credible intervals causes the graphs to become unreadable.

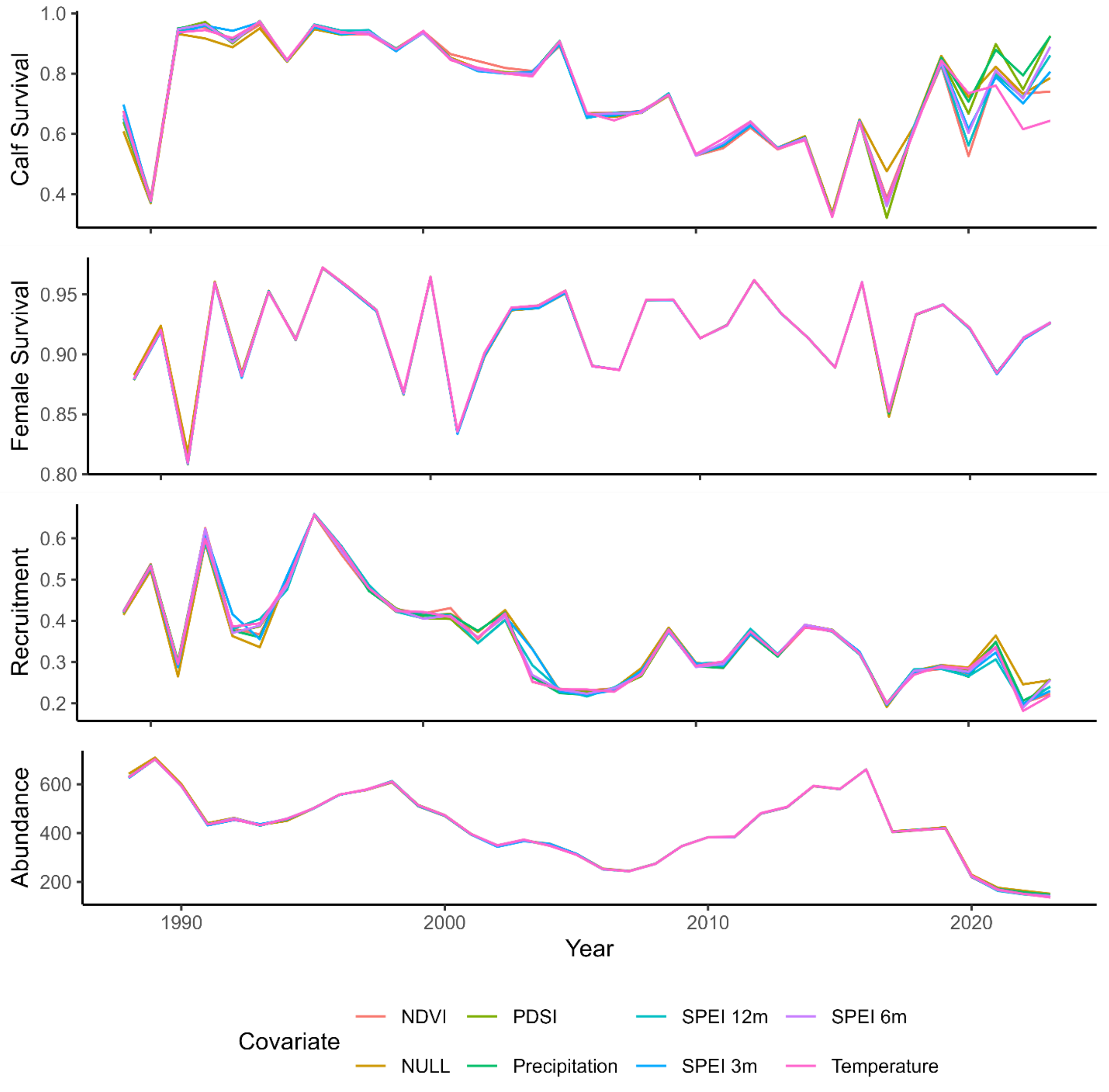


Figure S1.3

Slope estimates for the covariates on recruitment (top panel) and calf survival (bottom panel) from models using each of the 7 climate covariates. The points represent the medians and the lines represent the symmetrical 95% credible intervals.

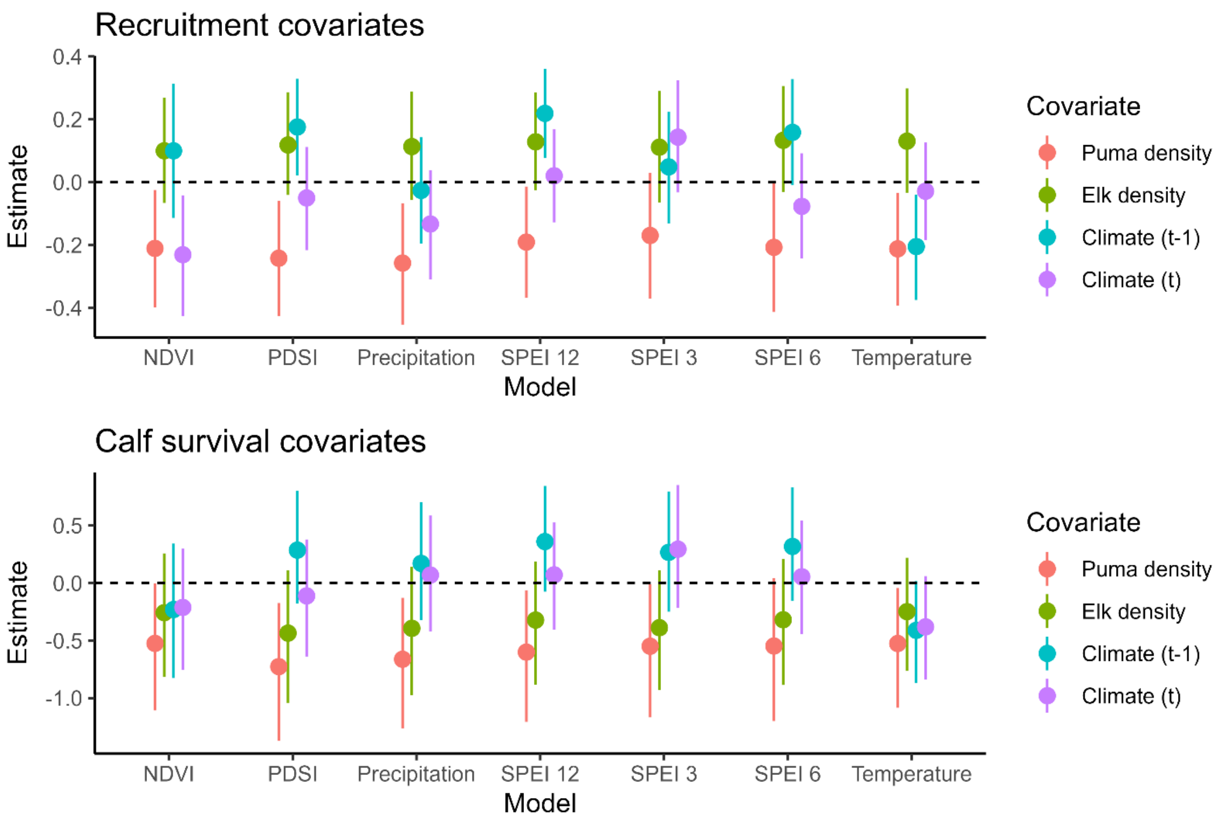
