## Appendix S2 - IPM Formulation for "The effects of predator recovery and climate change on the long-term demography of a flagship herbivore"

**SI 2: IPM Formulation**

Building a model that incorporates data from 5 sources spanning 36 years requires making choices and seemingly endless troubleshooting. This SI walks through some of those choices and troubleshooting milestones in more detail than fit into the main text. It also analyzes how the results from the version of the model we present compare to some of the potential alternative formulations.

*SI 2.1 – Removals*

One unique aspect of the data from Starkey is the "known integer" aspect of some of the data, *i.e.* the number of elk harvested from Starkey and added to or removed from Starkey are known exactly each year. Incorporating additions into a population process model is simple: just add that number to the appropriate age class at the corresponding time step. Removals, theoretically, work the same way but present a new issue: negative elk. Unlike when a fraction of a population dies or emigrates, a number being subtracted can result in more elk leaving that existed to begin with. This lead to numerous headaches, usually involving nodes that were inconsistent with parents. We found 3 choices vital to handling the removal data effectively: using normal approximations throughout the process model, truncating distributions at the minimum number of elk known, and setting the initial values well.

Using normal approximations means that negative elk are not impossible (nor are more elk surviving than existed), from a probability standpoint. The error of nodes not matching parents resulted from impossible values passed to the binomial distribution, such as a negative number of trials or the number of successes exceeding the number of trials. Using normal approximations to allow biological impossibilities to be possible from the probability standpoint bypassed errors and allowed the model to run. To prevent biological impossibilities, we leveraged the truncation functions.

In any given year, the elk population had a floor, defined by the number of individuals that were known alive either during or both before and after the year (ignoring temporary emigration is one perk of closed populations). We used that minimum population to truncate the potential outcomes of survival and recruitment. Because harvested elk were “observed” in the year they were harvested, they contributed to the minimum population in the prior years. With the inclusion of harvested elk in the minimum population, truncation did not only keep the population above 0; it also kept the population high enough to support the future years’ harvest. It also more fully incorporates our knowledge of the population to inform the estimates of vital rates.

Finally, initial values are critical in removal models. There most always be a large enough population to support the number of animals removed. We used relatively high (>0.9) initial values for survival and recruitment to ensure that, at the start of the MCMC, the numbers were possible, if unlikely. Another reasonable option would be to set the initial values based on the posteriors of a previous run, but that requires a prior successful run and the willingness to change initial values that worked. Theoretically, quality initial values alone would overcome the removals issue. However, we found that, after the burn-in phase, the mismatched node/parent issue would prevent the sampling phase from starting. Bypassing the burn-in and manually discarding some number of samples from the posterior as “burn-in” could bypass that problem.

*SI 2.2 – Samplers*

We used both nimble (NIMBLE Development Team, 2024) and JAGS (Plummer, 2003) for MCMC sampling because neither met all of our needs. We found JAGS to be extremely slow compared to nimble. The computer used for the bulk of the model runs has an Intel Core i7-10610U processor and 32GB of RAM. On that machine, the CJS model took days to complete 50,000 iterations (which was at least double the number needed) in JAGS and only ~12 hours in nimble. Unfortunately, the IPM consistently failed in nimble. The two sampler configurations that we tried worth noting are nimble's default sampler configuration (*e.g.* letting nimble set the samplers) and using the nimble option `onlySlice = T` to set all of the samplers to slice samplers. When using the default nimble sampler, the MCMC chains would not update; they would stay on the exact same value for millions of iterations, regardless of our choices for initial values. When using the slice samplers, the MCMC chains diverged wildly from reasonable state spaces, eventually estimating upwards of $10^8$ elk in 100 $km^2$. We suspect there is a sampler configuration that would allow the IPM to fit in nimble, but we did not find it. The IPM ran and converged well in JAGS (albeit over more hours than we would have liked), so we ran the component models in nimble and the IPM in JAGS.

*SI 2.3 – Partial Bayesian Approach*

Running the IPM in stages, with the component models being run first and summaries of those results being passed as data to the IPM (the partial Bayesian approach), gave tremendous speed gains, but might introduce bias. The speed gains came both from being able to use nimble for the component models and from being able to run the intensive CJS model for the number of iterations needed for it to converge rather than the (much higher) number of iterations for the IPM to converge. We tested the speed of a fully integrated version of the IPM (*i.e.* having all of the observation models connected through a process model within a single, integrated model) and found that, for a single chain to converge and explore the posterior, it would need to run for ~ 250 days, assuming the number of iterations required was the same as the partial Bayes approach. Nine months is a bit too long to wait for results troubleshooting (or to have a personal computer run reliably and continuously), so we ran smaller versions of the partial Bayes approach and the fully integrated model to assess whether they give similar results.

To test smaller versions of the models, we divided the study period into 6 smaller study periods (1988-1994; 1994-2000; 2000-2006; 2006-2012; 2012-2018; 2018-2023) and built the data for the IPM and each of the observation models using only information collected in those years. We then fit the two versions of the IPM to each set of data. Subsetting the data in this way reduced the size of the CJS data from 2708 by 36 to ~400 by 7 and allowed the fully integrated model to run in ~12 hours. We used 12-month SPEI as the climate covariate in all of the models and did not include elk density to avoid running null models prior to the test. In total, each model included 3 covariates: the climate covariate from year *t*, the climate covariate from year *t-1*, and puma density. Each covariate was included for both recruitment and calf survival, giving a total of 6 betas estimated for each test.

We found that the partial Bayes approach gave similar results to the fully integrated model with a few exceptions (Fig. S2.1). Across demographic rates, the fully integrated model tended to give more precise annual estimates than the partial Bayes approach, while the partial Bayes approach was slightly more precise in estimates of N (Fig. S2.1). Neither model estimated a parameter with a median falling outside of the other model’s 95% credible interval without underlying differences in the data.

The models diverged in two places due to underlying differences in the data. Most notably, the fully integrated model estimated dramatically lower female survival than the partial Bayes approach in 2017 and 2020. In both 2017 and 2020, intentional elk reductions occurred, with many elk being released from the winter feed grounds outside of Starkey. In that process, not all of the individual identities of the removed elk were recorded. In the CJS model, those capture histories are indistinguishable from the capture history of individuals that died between 2016/17 or 2019/20. They are also indistinguishable from individuals that lived past that year but were never captured again, but those capture histories become less likely with high capture probability and each additional year of data. Because we know the survival estimates from those years are artificially low, we withheld them from the main-text version of the IPM. For consistency we withheld them from the partial Bayes tests as well, but we did not program any exceptions for those years in the fully integrated tests. Those two survival estimates are spurious due to the data and likely widened the credible intervals of the survival estimates from 2013-2023 due to their inclusion in the random distribution of each survival parameter.

The second place the models diverge due to the structure of the test is in the final survival estimates of each time period (1994, 2000, 2006, 2012, 2018, and 2023). In the CJS component of the partial Bayes approach, both survival and detection probability are allowed to vary freely by year. In that formulation of the CJS model, only the product of survival and detection probability is estimable in the final year (Otis et al., 1978). Because survival is not estimable in those years, we withhold them from the partial Bayes tests and withheld 2023 survival estimates from the main text version of the IPM. Thus, in each final year of the partial Bayes tests, the only information informing survival is the random effect based on the five prior years and the count data, leading to particularly wide credible intervals for calf survival (Fig. S2.1). In contrast, in the fully integrated model, there are random effects on both survival and detection probability, allowing the CJS model to contribute to survival estimates and leading to more precise estimates of survival.

Most importantly to our analyses, the estimates of the effect of each covariate on both calf survival and recruitment were consistent between the fully integrated model and the partial Bayes approach (Fig. S2.2). The major deviations between the two models were all on survival parameters, which have slightly different underlying data as noted above. Additionally, when the precision of the estimates differed between models, the partial Bayes approach was almost always less precise than the fully integrated model, reducing the probability of type 1 errors.

Figure S2.1: Estimates of recruitment, calf survival, adult female survival, and total abundance each year from the fully integrated model and the partial Bayes approach of the IPM. Estimates arise from models fit to 7 years of data in sequence, rather than from the full sequence fit in a single model. In total, 6 time periods were used. The fully integrated model estimates lower survival in 2017 and 2020 due to differences in the underlying data (see SI text). Similarly, the survival estimates in every sixth year of the partial Bayes approach are less precise due to the structure of the underlying CJS model.

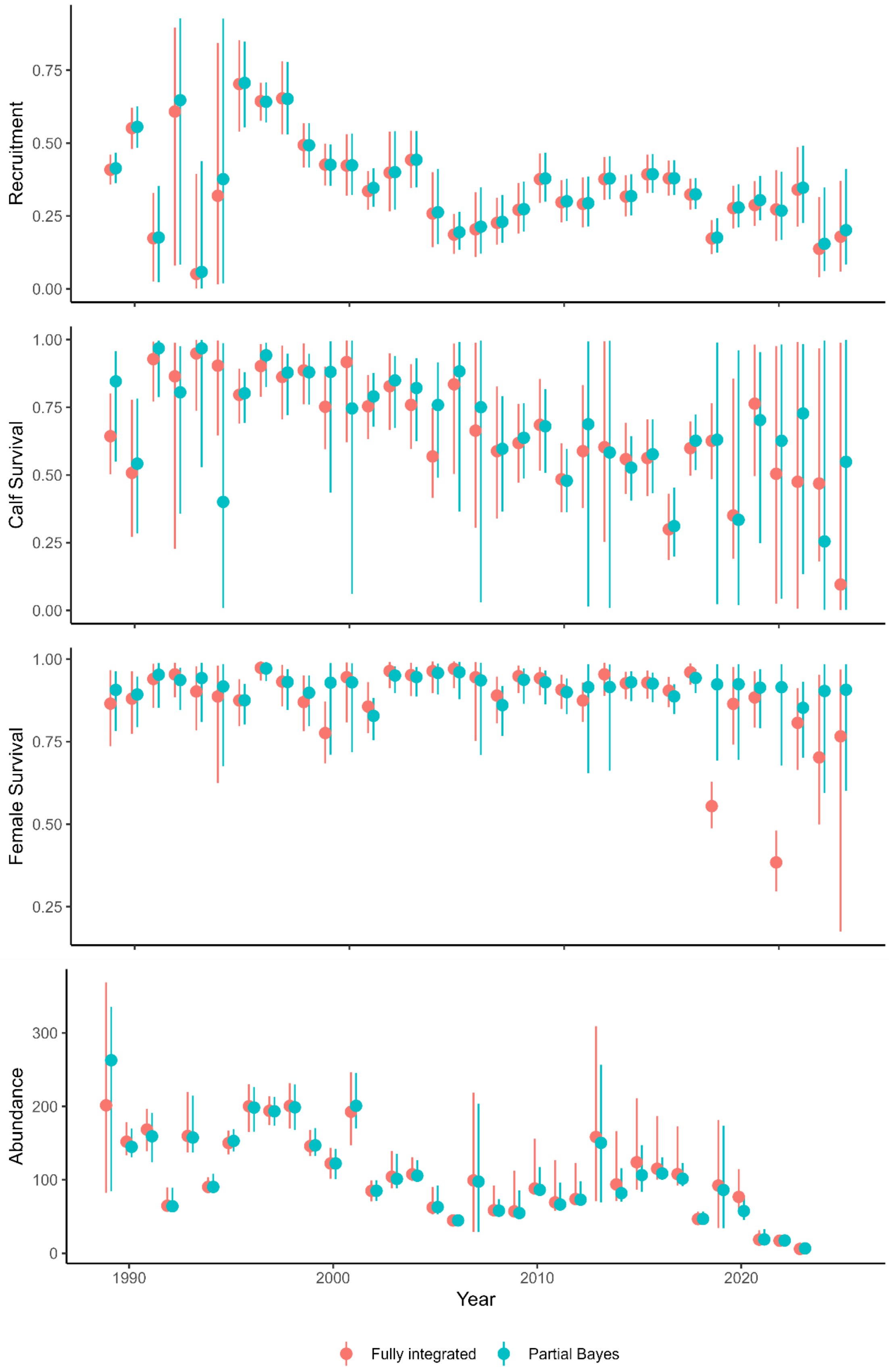
Recruitment
0.00
0.25
0.50
0.75
Calf Survival
0.00
0.25
0.50
0.75
1.00
Female Survival
0.25
0.50
0.75
1.00
Abundance
0
100
200
300
1990
2000
2010
2020
Year
Fully integrated
Partial Bayes

Figure S2.2: Scatter plots of the median (left panel) and variance (right panel) of the posterior distributions of the beta estimates from the partial Bayes approach (x-axis) against the same parameters from the fully integrated model (y-axis). Circles represent effects on calf survival, and triangles represent effects on recruitment. The color indicates the covariate.

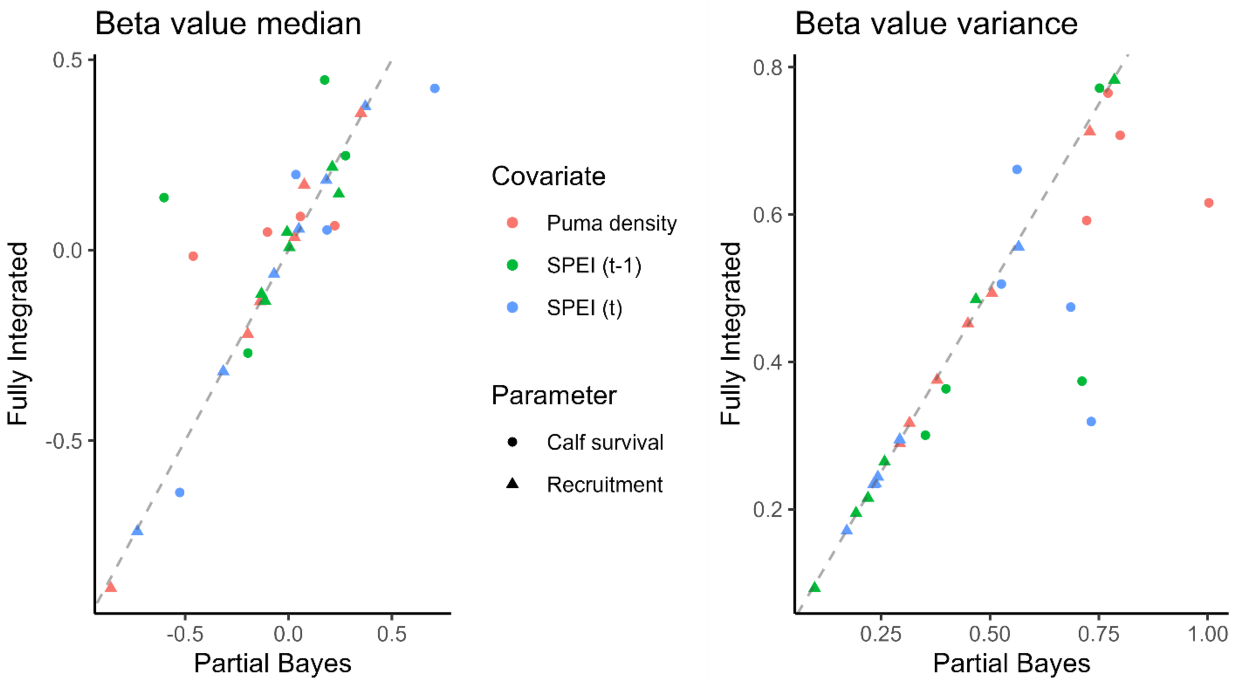

*SI 2.4 – Sensitivity to abundance data*

We evaluated two aspects of the incorporation of abundance information in the IPM for potential biases in the model. First, we tested whether an error structure for the corrected counts that more accurately reflected the count correction process would allow more variance than the normal error structure typically used in IPMs. To do that, we calculated alternative standard deviations in two ways. We first propagated the uncertainty around detection probability from the CJS model into the counts by dividing the naïve counts by the full detection probability posterior distribution. We then calculated the standard deviation of the count in each year as the standard deviation of the resulting counts.

The second approach we took to an error structure that reflected the process was calculating the standard deviation of the negative binomial distribution corresponding to the count data process. To do that, we used the formulation of the negative binomial that treats the number of trials necessary to record a given number of successes as a random variable. We treated the naïve count as the number of success and the latent abundance as the random variable. We used the median of the posterior distribution of detection probability for the probability parameter in the negative binomial. We then calculated the standard deviation following: SD = sqrt(r*(1-p)/p^2), where r is the naïve count and p is the detection probability.

We compared the two estimates of the standard deviation of counts (Fig. S2.3) to the estimated standard deviation of counts from the IPM. In all years, the standard deviations calculated from the detection process were much smaller than the estimated standard deviation using the normal error structure in the IPM. Because the normal error structure was less informative than the alternatives, we retained it in the final version of the IPM.

Figure S2.3 Estimated standard deviations of counts for each year from two methods. Posterior indicates that the standard deviation was calculated as a summary statistic of the naïve count divided by the posterior distribution of detection probability. NB stands for negative binomial and is the standard deviation of the random variable N – the number of elk in the population that results in the observed naïve count given the detection probability. It is calculated as SD = sqrt(r*(1-p)/p^2), where r is the naïve count and p is the detection probability. The median of the posterior distribution was used for p. For reference, the median estimated standard deviation of female counts in the IPM was ~95.

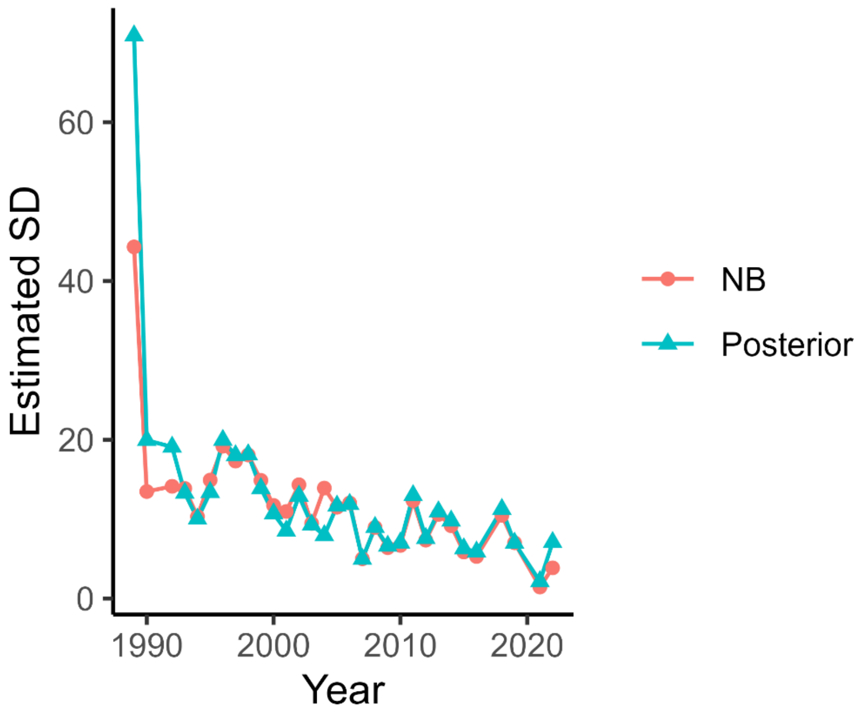


We also assessed whether the historic abundance estimates were having significant effects on the parameter estimates in the IPM. To do that, we ran a version of the IPM without the abundance estimates included. In that version, only the corrected counts directly informed abundance. We compared the estimates and precision of the demographic parameters (Fig. S2.4) and the beta values (Fig. S2.5) from the two versions of the IPM. We found no systematic differences in the relative values, so we retained the historic abundance estimates in the model to better anchor the IPM abundance estimates to the actual number of elk in Starkey throughout our study.

Fig. S2.4 Scatterplots of selected vital rates from the main text version of the IPM (x-axis) and the same version of the IPM with the abundances estimates excluded (meaning only the count data informed abundance). All of the vital rates were centered and scaled to facilitate simple linear regression. The black lines are the 1:1 line. The blue lines are the best fit lines and the shaded areas represent the uncertainty around the slope. Best fit lines and uncertainty were calculated using the `geom_smooth()` method in the R package ggplot2, with the method option set to `method = “lm”`. No best fit line deviates from the 1:1 line.

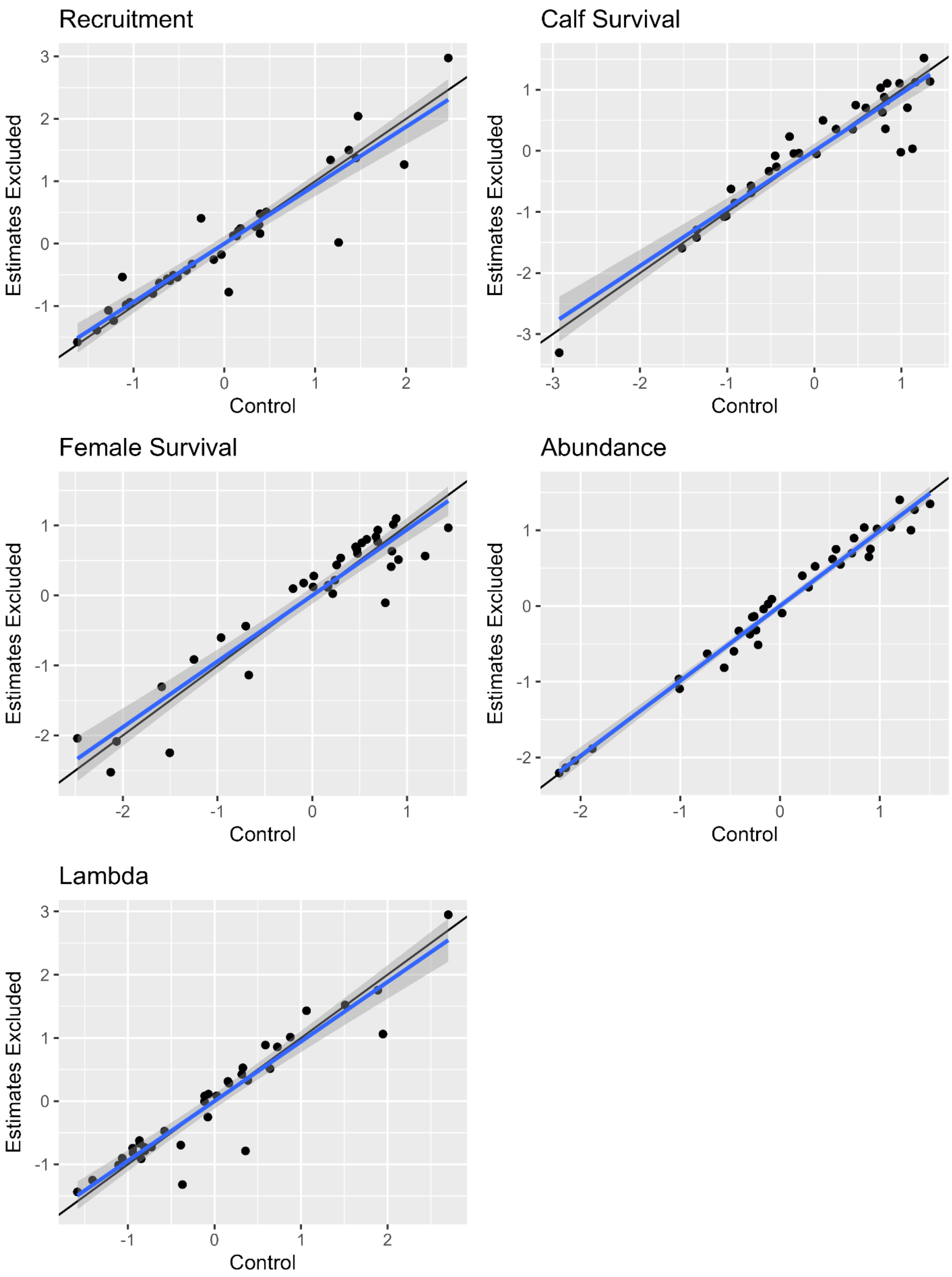
Recruitment
Calf Survival
Female Survival
Abundance
Lambda
Estimates Excluded
Control
3
2
1
0
-1
-2
-3

Fig. S2.5 Scatterplot of the beta estimates from the main text IPM (control, x-axis) and the IPM with the abundance estimates excluded (y-axis). The teal points are betas on recruitment and the red points are betas on survival. The dotted line is the 1:1 line.

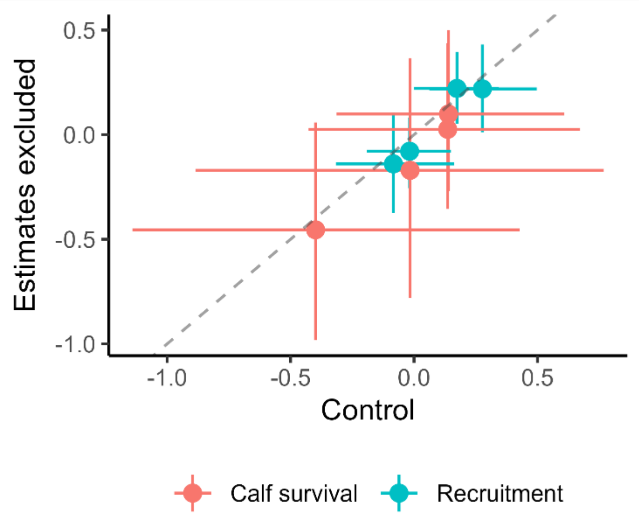
