## Appendix S3 - Residual Plots for "The effects of predator recovery and climate change on the long-term demography of a flagship herbivore"

SI 3 – Residual plots and additional marginal plots

Figure S3.1: Residuals from the recruitment component of the IPM plotted against (a) the puma density index, (b) female abundance, (c) SPEI of the current year, and (d) SPEI of the prior year. Residuals were calculated as the difference between estimated recruitment and recruitment predicted by the logistic model and the posteriors of the covariate betas, meaning each residual is represented by a posterior distribution. The dots represent the median and the vertical lines the 95% credible intervals of the residuals.

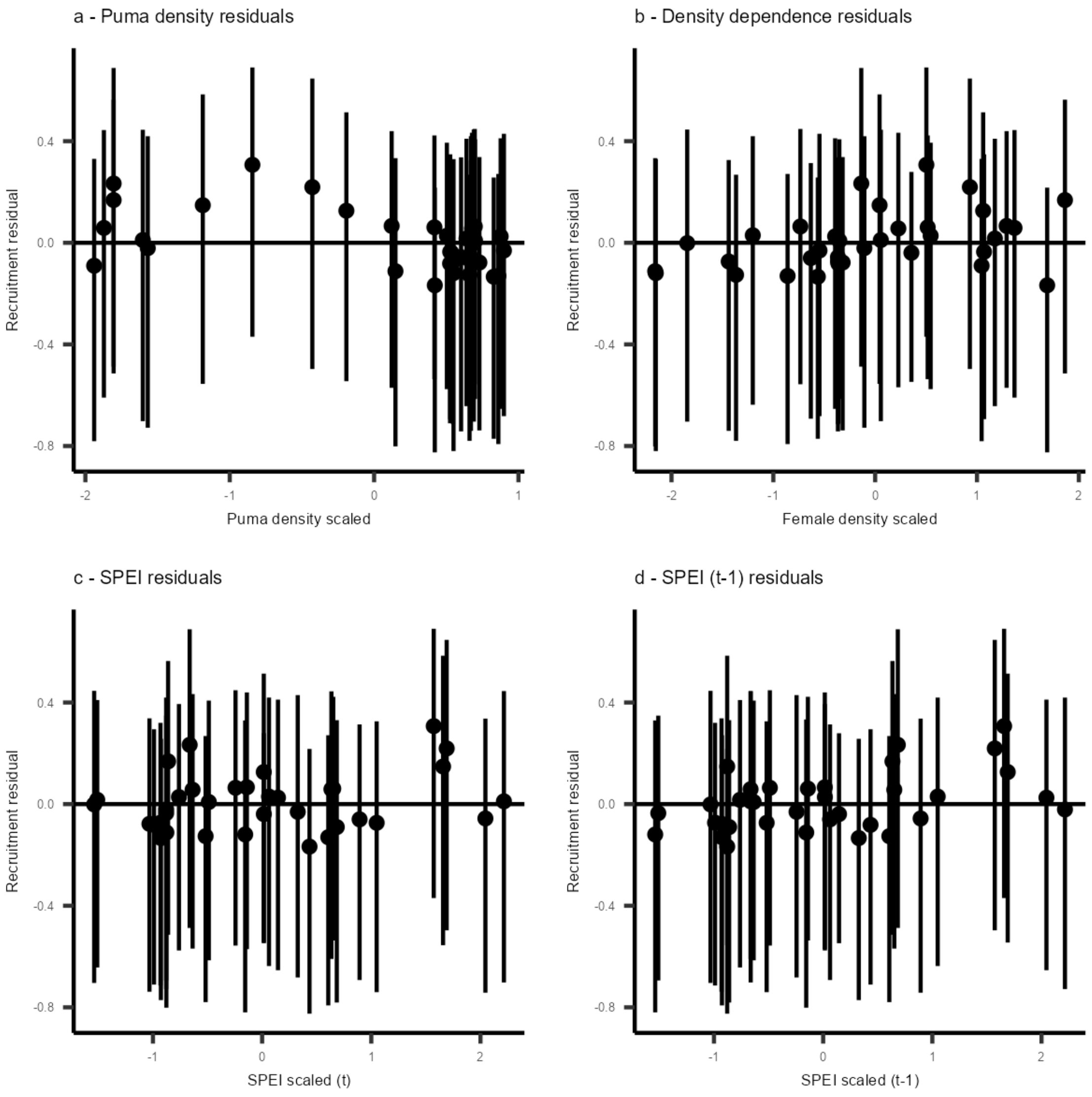

Figure S3.2 Residuals from the survival component of the IPM plotted against (a) the puma density index, (b) female abundance, (c) SPEI of the current year, and (d) SPEI of the prior year. Residuals were calculated as the difference between estimated recruitment and recruitment predicted by the logistic model and the posteriors of the covariate betas, meaning each residual is represented by a posterior distribution. The dots represent the median and the vertical lines the 95% credible intervals of the residuals.

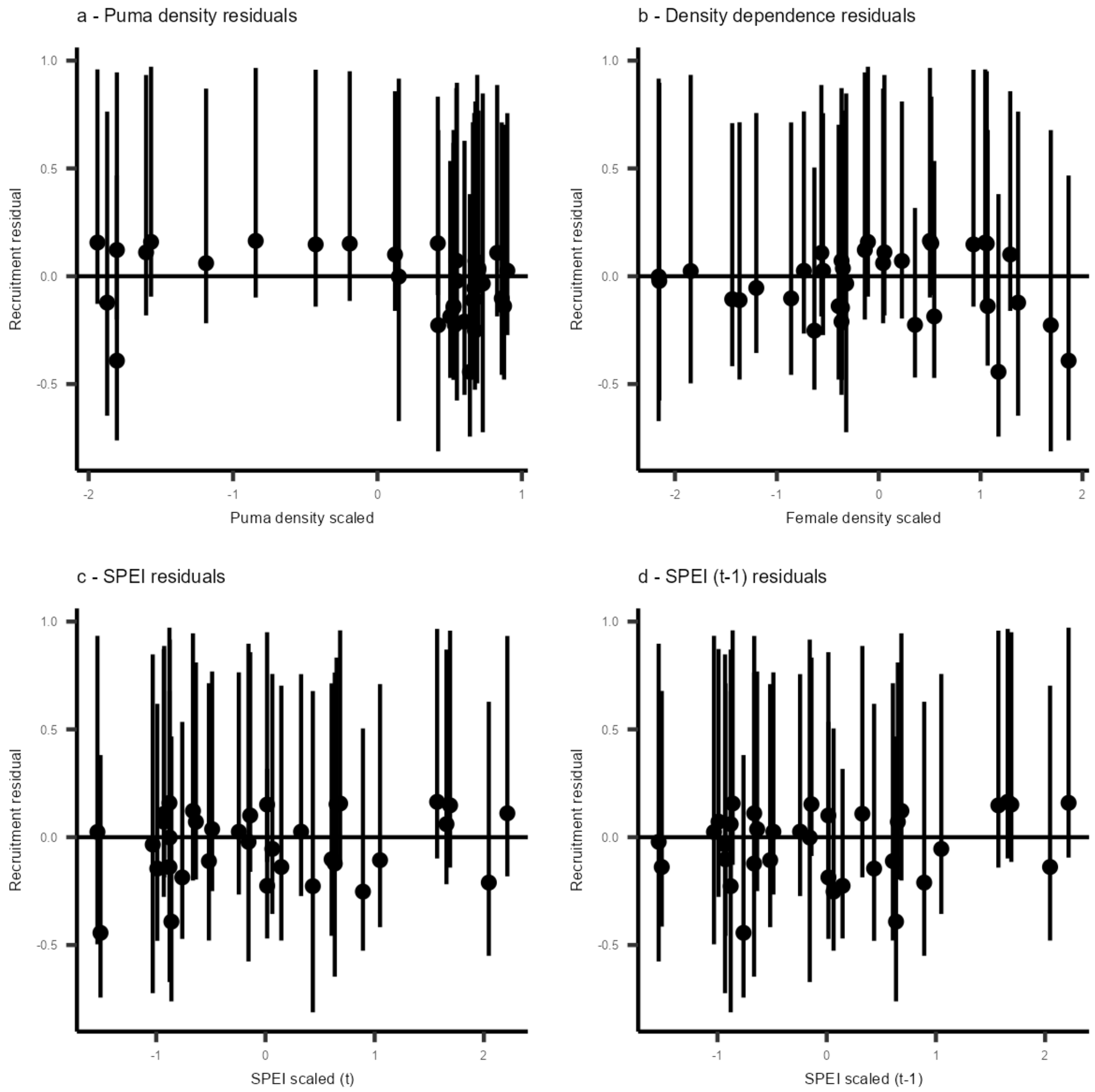

Figure S3.3 and S3.4: Residuals from the pregnancy rate models. Each point represents the median difference between estimated pregnancy rate and the predicted value of pregnancy rates from the model. The bars represent 95% credible intervals. Residuals are separated by age/sex class and by covariate (elk density and SPEI).

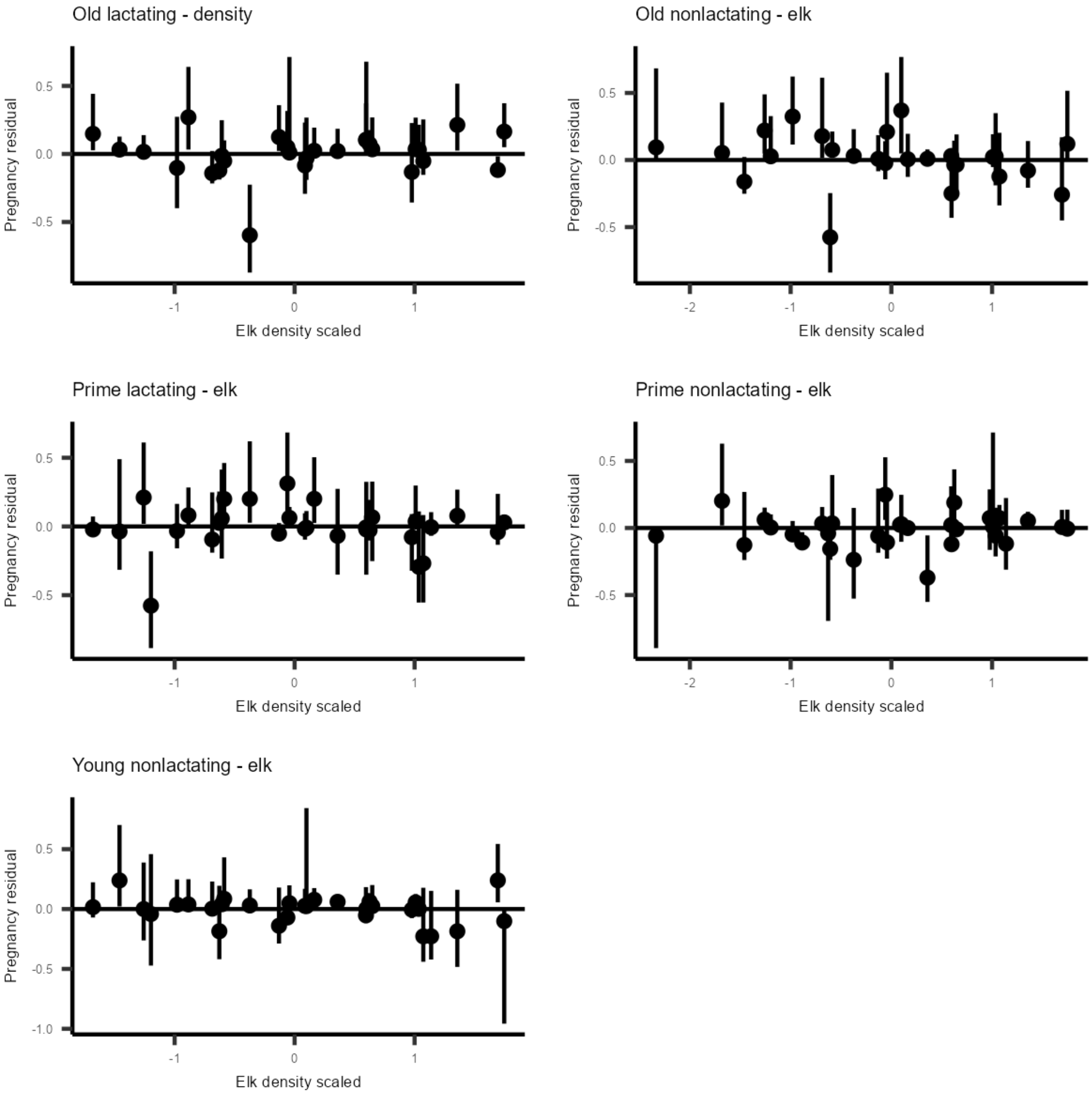

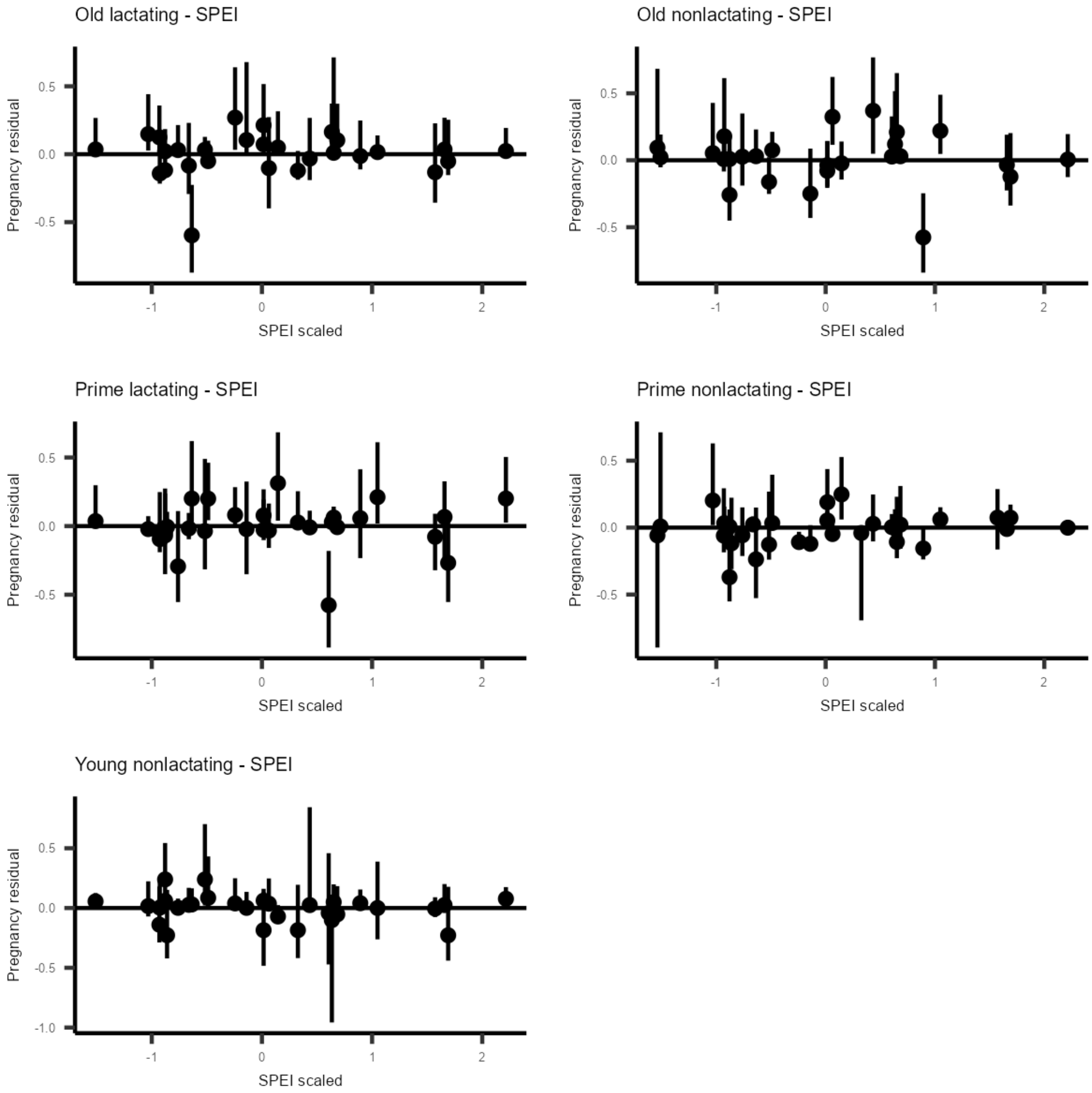
Old lactating - SPEI
Pregnancy residual
0.5
0.0
-0.5
-1
0
1
2
SPEI scaled
Old nonlactating - SPEI
Pregnancy residual
0.5
0.0
-0.5
-1
0
1
2
SPEI scaled
Prime lactating - SPEI
Pregnancy residual
0.5
0.0
-0.5
-1
0
1
2
SPEI scaled
Prime nonlactating - SPEI
Pregnancy residual
0.5
0.0
-0.5
-1
0
1
2
SPEI scaled
Young nonlactating - SPEI
Pregnancy residual
0.5
0.0
-0.5
-1.0
-1
0
1
2
SPEI scaled

Figure S3.5: Marginal plots for the pregnancy models of prime-aged, nonlactating females, old females, and young females (see the main text for prime-aged, lactating females). The left column is marginal plots with respect to elk density and the right column is marginal plots with respect to SPEI. The line represents the median predicted values, and the shaded area represents 95% credible intervals. The points are the median estimates of pregnancy rates, and the bars are 95% credible intervals.

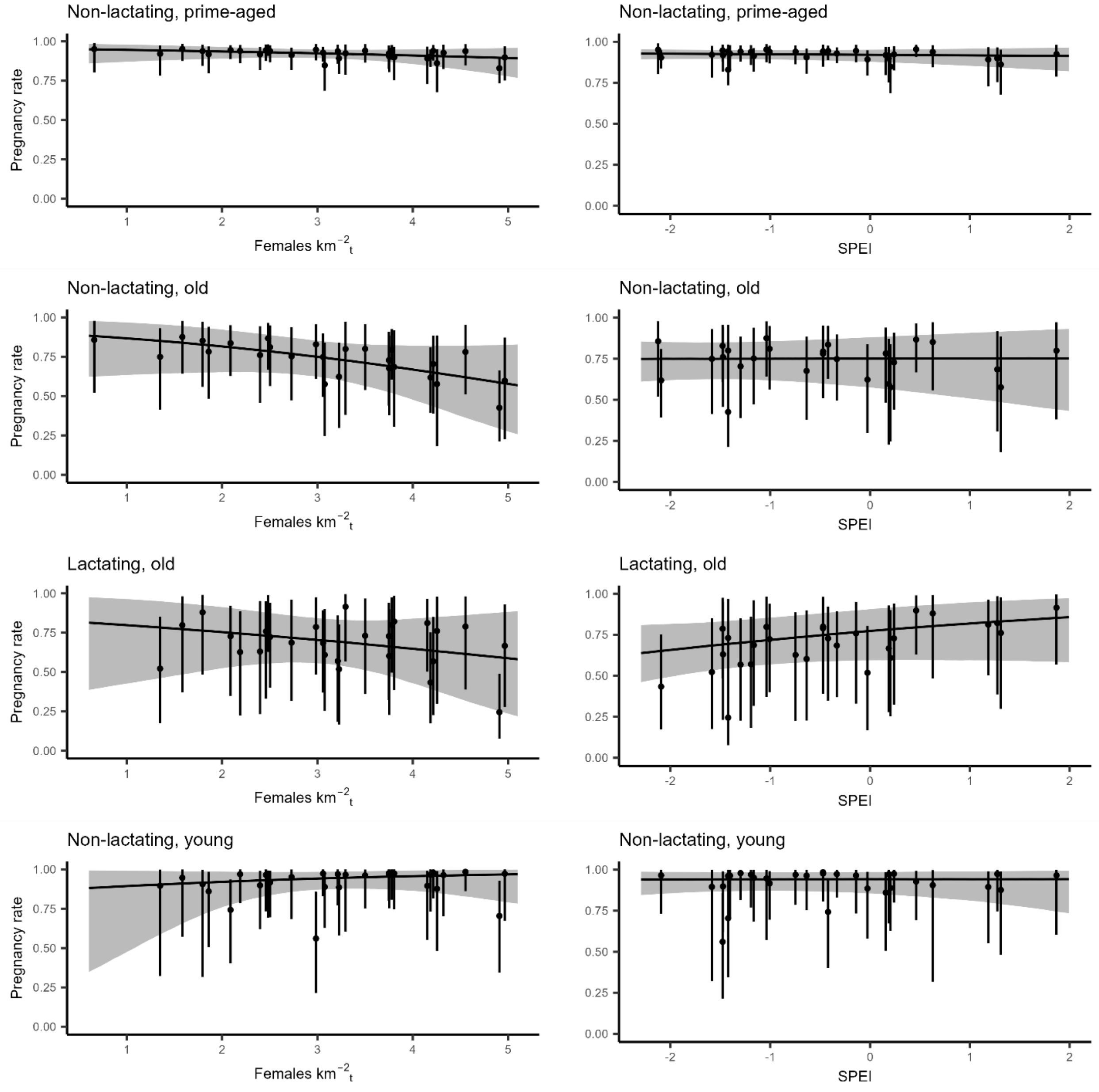
